# A Rab6 to Rab11 transition is required for dense-core granule and exosome biogenesis in *Drosophila* secondary cells

**DOI:** 10.1101/2023.04.04.535541

**Authors:** Adam Wells, Cláudia C. Mendes, Felix Castellanos, Phoebe Mountain, Tia Wright, S. Mark Wainwright, M. Irina Stefana, Adrian L. Harris, Deborah C. I. Goberdhan, Clive Wilson

## Abstract

Secretory cells in glands and the nervous system frequently package and store proteins destined for regulated secretion in dense-core granules (DCGs), which disperse when released from the cell surface. Despite the relevance of this dynamic process to diseases such as diabetes and human neurodegenerative disorders, our mechanistic understanding is relatively limited, because of the lack of good cell models to follow the nanoscale events involved. Here, we employ the prostate-like secondary cells (SCs) of the *Drosophila* male accessory gland to dissect the cell biology and genetics of DCG biogenesis. These cells contain unusually enlarged DCGs, which are assembled in compartments that also form secreted nanovesicles called exosomes. We demonstrate that known conserved regulators of DCG biogenesis, including the small G-protein Arf1 and the coatomer complex AP-1, play key roles in making SC DCGs. Using real-time imaging, we find that the aggregation events driving DCG biogenesis are accompanied by a change in the membrane associated small Rab GTPases which are major regulators of membrane and protein trafficking in the secretory and endosomal systems. Indeed, a transition from *trans*-Golgi Rab6 to recycling endosomal protein Rab11, which requires conserved DCG regulators like AP-1, is essential for DCG and exosome biogenesis. Our data allow us to develop a model for DCG biogenesis that brings together several previously disparate observations concerning this process and highlights the importance of communication between the secretory and endosomal systems in controlling regulated secretion.

**Author summary:** Cells communicate with each other by releasing signalling molecules that bind receptors on target cells and alter their behaviour. Before their release, these signals are typically stored in condensed structures called dense-core granules (DCGs). DCGs are found in many animal species and their dysregulation is linked to several major human diseases, such as diabetes and neurodegenerative disorders. However, the mechanisms controlling DCG formation and secretion are only partly understood. Here we study this process in fruit flies using a secretory cell, which contains unusually large DCGs. We show that known regulators of DCG formation in mammals also control DCG production in these fly cells and identify new DCG assembly steps by following the process in living cells. Furthermore, we show that the cell’s secretory and recycling endosomal compartments must interact to induce the rapid condensation of proteins into a DCG, and that known regulators of DCG formation are needed for this crucial event to take place. Our work provides a platform from which to work out the molecular mechanisms that enable this critical secretory-endosomal interaction and probe its roles in diseases of secretion.

## Introduction

Proteins are secreted from eukaryotic cells by both constitutive and regulated pathways. Hormone-, enzyme- and neuropeptide-secreting cells, including endocrine adrenal chromaffin cells and pancreatic β cells, exocrine pancreatic acinar cells and neurons respectively, are typically specialised for regulated secretion. They package and store the proteins, which they will release, into dense-core granules (DCGs) within so-called dense-core vesicles (Gondré-Lewis et al., 2012).

By studying the DCG packaging process in cell lines that represent some of these different secretory cell types (e.g. Haeberlé et al., 2015; Tsuchiya et al., 2010; Giudici et al., 1992), several mechanisms controlling DCG biogenesis and release have been identified and shown to be shared between different cells. For example, clustering of cargos into immature DCG compartments in the *trans*-Golgi network requires cholesterol and lipid raft-like structures (Tsuchiya et al., 2010), together with specific enzymes that regulate lipid metabolism, namely phospholipase D1 and diacylglycerol kinase (Siddhanta et al., 2000). Granin proteins may be required for compartment budding and DCG assembly (Day and Gorr, 2003; Gondré-Lewis et al., 2012). Several cytosolic adaptor proteins, such as the AP-1 coatomer complex and Golgi-localising, γ-adaptin ear homology domain, ARF-binding proteins (GGAs) are recruited to the *trans*-Golgi by the small G protein Arf1 (Stamnes et al., 1993; Traub et al., 1993), and are then involved in trafficking molecules to and from maturing DCG compartments (Bonnemaison et al., 2013). The DCG maturation process is also dependent on reduced pH (Wu et al., 2001) and an increase in intraluminal calcium (Ca^2+^) ions (Yoo and Albanesi, 1990). However, the molecular and membrane trafficking processes that drive and coordinate these changes remain unclear. Once matured, the DCG compartments fuse to the plasma membrane via mechanisms requiring Ca^2+^- dependent synaptotagmins and vesicle-associated membrane proteins (VAMPs) (Messenger et al., 2014).

Rab GTPases are another key set of molecules involved in regulated secretion. These small monomeric GTPases regulate membrane trafficking and organelle identity in the secretory and endolysosomal systems (Barr, 2013). Analysis in mammalian cell lines has highlighted a role for Rab6 in the formation of immature DCG compartments at the *trans*-Golgi network (Miserey-Lenkei et al., 2010). However, mature dense-core vesicles have been reported to be associated with endosomal Rabs, like Rab11 (Sugawara et al., 2009), suggesting a potential cross-talk between the secretory and endosomal systems in regulated secretion.

Dissecting out the molecular mechanisms underlying regulated secretion is not only of biological importance, but of clinical relevance. For example, Type 1 and Type 2 diabetes involves aberrant secretory regulation in β cells (Cai et al., 2011; Carrat et al., 2020), while defective processing and secretion of specific proteins, such as the Amyloid Precursor Protein (APP), is associated with neurodegenerative diseases, including Alzheimer’s Disease (Arbo et al., 2020; Kuo et al., 2021). In mammals, however, cell biological studies of regulated secretion are usually undertaken in cultured cells, either isolated primary cells or immortalised cell lines, where it is difficult to reproduce the physiological microenvironment in which secretory cells function in living organisms. Furthermore, it is challenging to follow the events involved, because high-resolution imaging of DCG compartments, each of which is typically 0.2-1 µm in diameter, traditionally requires electron microscopy on fixed tissue.

The larval salivary gland (Burgess et al., 2012; Torres et al., 2014) and secretory cells of the proventriculus (Zhang et al., 2014) in the fruit fly, *Drosophila melanogaster*, as well as neurons and neuromuscular junctions in flies (James et al., 2014) and the nematode, *Caenorhabditis elegans* (Ailion et al., 2014), have provided *in vivo* genetic systems to study DCG formation and release in invertebrates, with several novel conserved regulators, some linked to endosomal trafficking, identified in *C. elegans* (e.g. Topalidou et al., 2016; Cattin-Ortolá et al., 2017). Generally, the DCG compartments in these cells are small and their substructures difficult to resolve with light microscopy. However, salivary gland granules increase in size during maturation, and it has therefore been possible to follow and genetically dissect some steps in this process (Ma et al., 2020). Such studies have, for example, shown that Rab1 and Rab6 localise to the membranes of maturing secretory granules, while Rab11 and Rab1 drive granule growth and maturation via mechanisms that are yet to be characterised (Ma and Brill, 2021).

We have characterised another cell system in *Drosophila*, the secondary cell (SC) of the male accessory gland, as a genetic cell model for regulated secretion (Fig. 1A, B, B’; Corrigan et al., 2014; Redhai et al., 2016; Fan et al., 2020; Marie et al., 2023). Its secretory compartments are exceptionally large, approximately 5 µm in diameter, making it possible to visualise the structures formed within. These include a ∼3 µm diameter DCG (Redhai et al., 2016), surrounded by intraluminal vesicles (ILVs), which are released as exosomes upon compartment fusion with the plasma membrane (Corrigan et al., 2013; Fan et al., 2020). ILV formation in SCs requires the core endosomal complexes required for transport (ESCRT) proteins as well as specific accessory ESCRT proteins, some of which also affect DCG biogenesis (Marie et al., 2023). Some immature secretory compartments, which lack DCGs, are marked by Rab6, but all mature compartments carry Rab11 at their surface (Prince et al., 2019; Fan et al., 2020), further supporting a link between endosomal and secretory pathways in DCG biogenesis. Although only a small subset of ILVs formed in these latter compartments are labelled by Rab11, the entire population of exosomes produced by these compartments are termed Rab11-exosomes (Fan et al., 2020). The Rab11- exosome pathway and its regulation appear to be conserved from fly to human cells and Rab11-exosomes have important physiological and pathological functions (Marie et al., 2023), making the further characterisation of this trafficking route of considerable interest.

**Fig 1.**
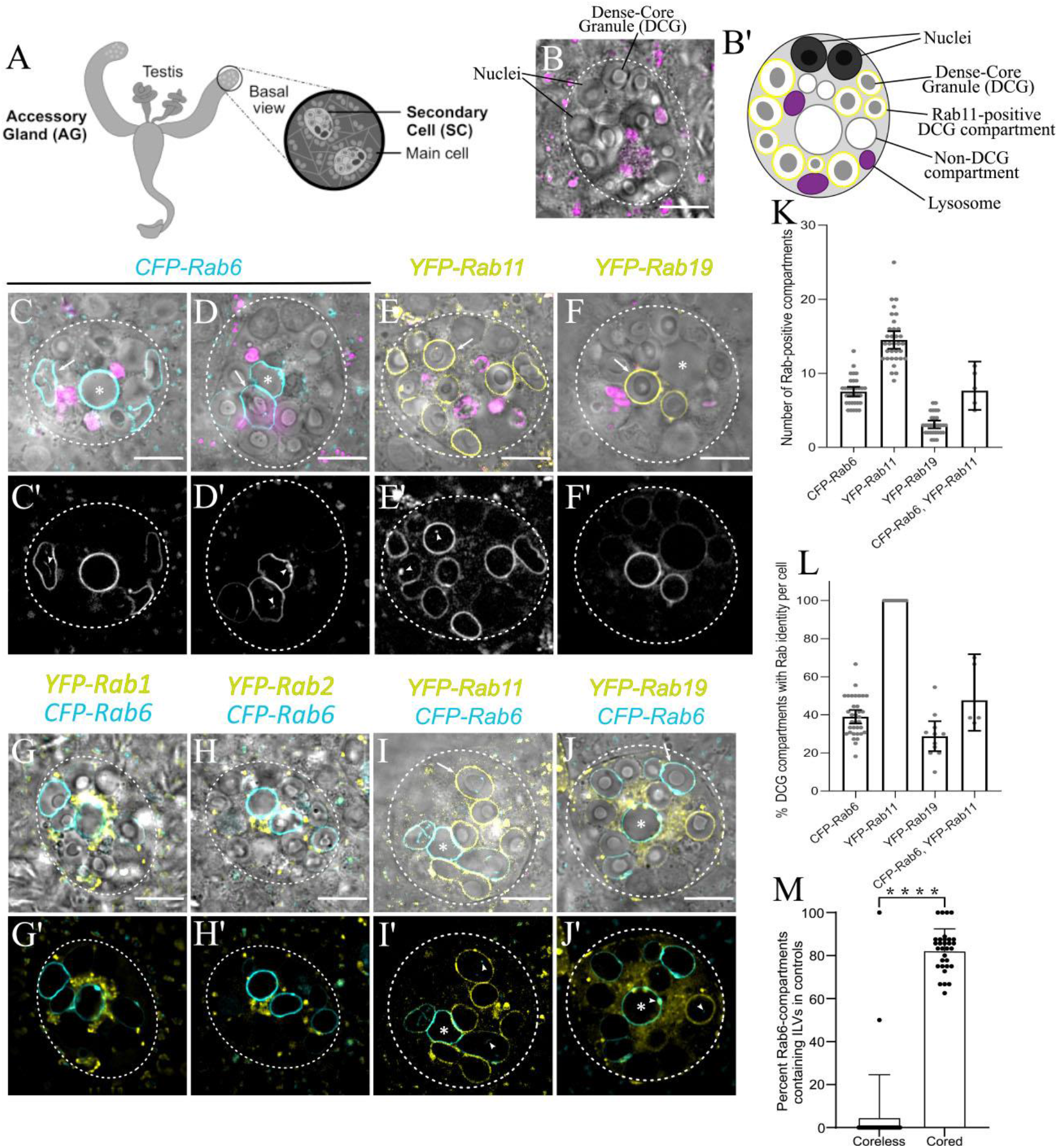
Morphology and Rab identity of DCG compartments in *Drosophila* secondary cells (A) Schematic of *Drosophila* accessory gland (AG), showing secondary cells (SCs) at the distal tip of each lobe. (B) *Ex vivo* Differential Interference Contrast (DIC) image of an SC from the AG of a six-day-old virgin male fly, stained with LysoTracker Red. (B’) Schematic of SC, with equivalent structures labelled. (C-J) DIC images of SCs, overlaid with LysoTracker Red (C-F), and fluorescent signal from different *Rab* gene traps (C-J). (C’-J’) Fluorescence-only images showing expression of each Rab. Arrowheads indicate ILVs labelled by various Rabs, which lie inside compartments. (C, D) CFP-Rab6-labelled compartments have a range of morphologies including spherical non-DCG compartments (C, *), irregularly shaped, non-DCG compartments (D, *), and DCG-containing compartments (arrows). (E) YFP-Rab11 marks all DCG compartments. (F) YFP-Rab19 marks two or three DCG compartments. (G, H and G’, H’) Small YFP-Rab1- (G) and YFP-Rab2-positive (H) clustered compartments surround central non-DCG-containing, CFP-Rab6 compartments. (I) Some YFP- Rab11-positive DCG compartments are labelled with CFP-Rab6. CFP-Rab6 puncta are also observed inside some compartments that are not Rab6-positive (arrow) and in a compartment that is Rab6- and Rab11-positive, but lacks a DCG (I, *). (J) YFP- Rab19 compartments are not co-labelled with CFP-Rab6. However, YFP-Rab19 and CFP-Rab6 do co-label microdomains and internal membranes on Rab6-marked compartments, eg. arrowhead in centre (J’), while Rab6-positive internal puncta are also found inside Rab19-positive compartments (arrowhead on right). (K) Histogram showing number of large compartments positive for different fluorescent Rabs in individual SCs. (L) Histogram showing % of DCG compartments labelled with specific fluorescent Rabs in individual SCs. (M) Histogram showing the proportion of CFP- Rab6-positive DCG compartments and coreless CFP-Rab6 compartments containing Rab6-positive ILVs in individual SCs expressing a control *rosy*-RNAi. Data for histograms were collected from three SCs per gland derived from 10 glands, except for the genotype expressing CFP-Rab6 and YFP-Rab11, where the relative expression levels of both fusion proteins varied considerably between different cells, so only some cells were suitable for analysis. Approximate outlines of SCs are marked by dashed circles. Scale bars: 10 µm. * marks representative non-acidic compartments that lack a DCG. Arrows mark representative DCG compartments labelled by various Rabs. For M, bars show mean ± SD; P<0.0001: ****.

Here, employing genetic knockdown techniques and high-resolution imaging in SCs, we demonstrate several parallels between *Drosophila* SCs and mammalian cells in the regulation of DCG biogenesis. Furthermore, by visualising the process of DCG formation in real-time, we show that it is accompanied by a Rab6 to Rab11 transition, and not by trafficking of cargos from Rab6-positive to Rab11-positive compartments. This Rab transition, which is controlled by known regulators of mammalian DCG biogenesis, plays a critical role in DCG as well as Rab11-exosome biogenesis, presumably by establishing a DCG-inducing microenvironment inside these compartments.

## Results

### Rab6, Rab11 and Rab19 mark specific compartment subsets in the SC secretory pathway

Previous expression analysis, primarily in fixed SCs, using *Rab* gene traps, where a YFP-Rab fusion protein is produced at normal levels from the endogenous *Rab* locus (Prince et al., 2019), confirmed that DCG compartments are marked by Rab11 (Redhai et al., 2016), but also revealed that some secretory compartments are labelled by Rab6 and by Rab19, a poorly characterised Rab proposed to be involved in apical secretion (Dunst et al., 2015). To assess the overlaps in expression of these different Rabs in more detail, we employed a live imaging approach, where the morphology of SCs is much better preserved. The *Rab* gene traps, which express either a CFP- or YFP-Rab fusion protein, had no effect on the number of DCG compartments in SCs, as scored using differential interference contrast (DIC) microscopy (Fig. 1B; Supp. Fig. S1A).

SCs contained 11 ± 2.2 (mean ± SD) DCG compartments in SCs (n=75), none of which were stained by the acidic dye LysoTracker Red (eg. Fig. 1C-F; Supp. Fig. S1B, C), and an additional 2.7 ± 2.0 (n=34) large non-acidic compartments that did not contain a DCG. A CFP-tagged form of Rab6 localised to 7.2 ± 1.9 (n=34) of these non-acidic compartments, including all of those that lacked a DCG (Fig. 1C, D, K). Some of these latter compartments were enlarged and/or irregular in shape, though frequently, at least one spherical non-DCG compartment of similar size to the DCG compartments was positioned centrally within the SC (Fig. 1C).

Rab1 and Rab2 are two Golgi-associated Rabs that have been implicated in trafficking processes that take place around the *trans*-Golgi network and precede formation of DCGs. In yeast, Rab1-labelled Golgi membranes convert to Rab6-coated membranes at the *trans*-Golgi (Thomas et al., 2021), while in nematodes, Rab2-interacting proteins associated with the Golgi apparatus are required for DCG compartment formation (Ailion et al., 2014; Cattin-Ortolá et al., 2017). Both YFP-Rab1 (Fig. 1G) and YFP- Rab2 (Fig. 1H) fusion proteins expressed from gene traps were concentrated in Golgi cisternae around the Rab6-positive, non-DCG-containing central compartments of SCs, consistent with these latter compartments being generated at the surface of the *trans*-Golgi.

As previously reported (Fan et al., 2020), YFP-Rab11 marked all the large DCG-containing compartments in SCs (Fig. 1E, L). About 39.0 ± 10.1% of these were also marked by Rab6 (Fig. 1I, K, L). Occasionally, one large non-acidic compartment that lacked a DCG was also weakly labelled with Rab11 and it often contained internal CFP-Rab6 puncta (Fig. 1I). Based on our previous findings with the *YFP-Rab11* gene trap (Fan et al., 2020), this Rab6 is presumably packaged inside ILVs. CFP-Rab6 puncta were also observed in some DCG compartments that lacked CFP-Rab6 at their surface (Fig. 1I; Supp. Fig. S1D).

ILVs marked by CFP-Rab6 and/or YFP-Rab11 were often seen adjacent to the boundaries of DCGs (Supp. Fig. S1B) and in some cases, extended along an arc around the DCG surface (Supp. Fig. S1C). It was clear from fluorescence imaging that Rab-labelled ILVs can also form continuous bridge-like structures running from the compartment’s limiting membrane to the DCG, sometimes across a gap of one to two micrometres (Supp. Fig. S1B, S1C), an observation previously reported using membrane-associated exosome markers in SCs with more ubiquitous ILV-labelling profiles (Fan et al., 2020, Dar et al., 2021). Although the significance of this association remains unclear, it is notable that ILV puncta appeared almost exclusively in compartments with DCGs. In SCs, over 80% of CFP-Rab6-labelled compartments with DCGs also contained Rab6-positive ILVs (Fig. 1M). By contrast, only a few core-less Rab6-labelled compartments contained ILVs, with the vast majority of cells containing only ILV-deficient, core-less compartments.

Finally, YFP-Rab19 labelled the entire surface of 3.7 ± 1.3 large compartments (n=30), all of which contained DCGs (Fig. 1F, K, L). These compartments were never marked by CFP-Rab6 (Fig. 1J), although YFP-Rab19 did mark microdomains on the surface of other non-acidic compartments, including those labelled by CFP-Rab6 (Fig. 1J, Supp. Fig. S1E). In addition to these microdomains on compartment surfaces, YFP- Rab19 also localised internally within various compartments, including those not marked by Rab19 on their surface (Fig. 1J’). This indicates that Rab19 can be incorporated into specific populations of ILVs, perhaps through invagination of the microdomains we observe on the compartment surface.

In summary, Rab6, Rab11 and Rab19 mark the highly enlarged secretory compartments found in SCs. Rab6 labels all large non-acidic compartments that do not contain DCGs, whilst Rab11 marks all the compartments that do. There is also significant overlap between these two markers, with Rab6 and Rab11 often colocalising on DCG compartments and on sporadic non-DCG compartments. Rab19 marks a subset of DCG compartments, which is distinct from the subset of Rab6- labelled compartments.

### Small Rab1-positive compartments give rise to Rab6-positive endosomes

To begin to understand the dynamics of secretory compartment biogenesis in SCs, it was first important to determine how the various compartments we had identified related to each other. We therefore began by investigating the relationship between the small Rab1-positive compartments that contact larger Rab6-positive compartments (Fig. 1G). Since Rab1 is frequently associated with Golgi stacks, whilst Rab6 is often located and functions in the *trans*-Golgi network (Miserey-Lenkei et al., 2010), we hypothesised that the Rab1- and Rab6-labelled compartments in SCs might interact, either through trafficking between these compartment types or by Rab1- positive compartments maturing into Rab6-positive compartments, as observed in yeast (Thomas et al., 2021). To test this, we carried out time-lapse imaging of live SCs expressing the *YFP-Rab1* and *CFP-Rab6* gene traps.

Through these experiments, it became clear that Rab1-positive compartments directly transition into Rab6-positive ones as they mature (Fig. 2, Movie S1). During this process, a small Rab1-labelled compartment increases in size from approximately 0.5 μm in diameter (Fig. 2A-B), until it has reached between 2.5 μm and 10 μm in diameter. Concurrently with this dramatic increase in size, the strong labelling by Rab1 is gradually lost and the compartment accumulates Rab6 on its limiting membrane to form a large Rab6-positive compartment that lacks a DCG (Fig. 2D). This process takes between 30 and 60 minutes, and soon after its completion, the Rab6-positive compartment produced contracts in volume again, after which it can form a DCG (Fig. 2E). Therefore, time-lapse experiments not only demonstrate that Rab1-positive compartments are the direct precursors of Rab6-positive compartments, but also that they are the same compartments that will eventually go on to form DCGs.

**Fig 2.**
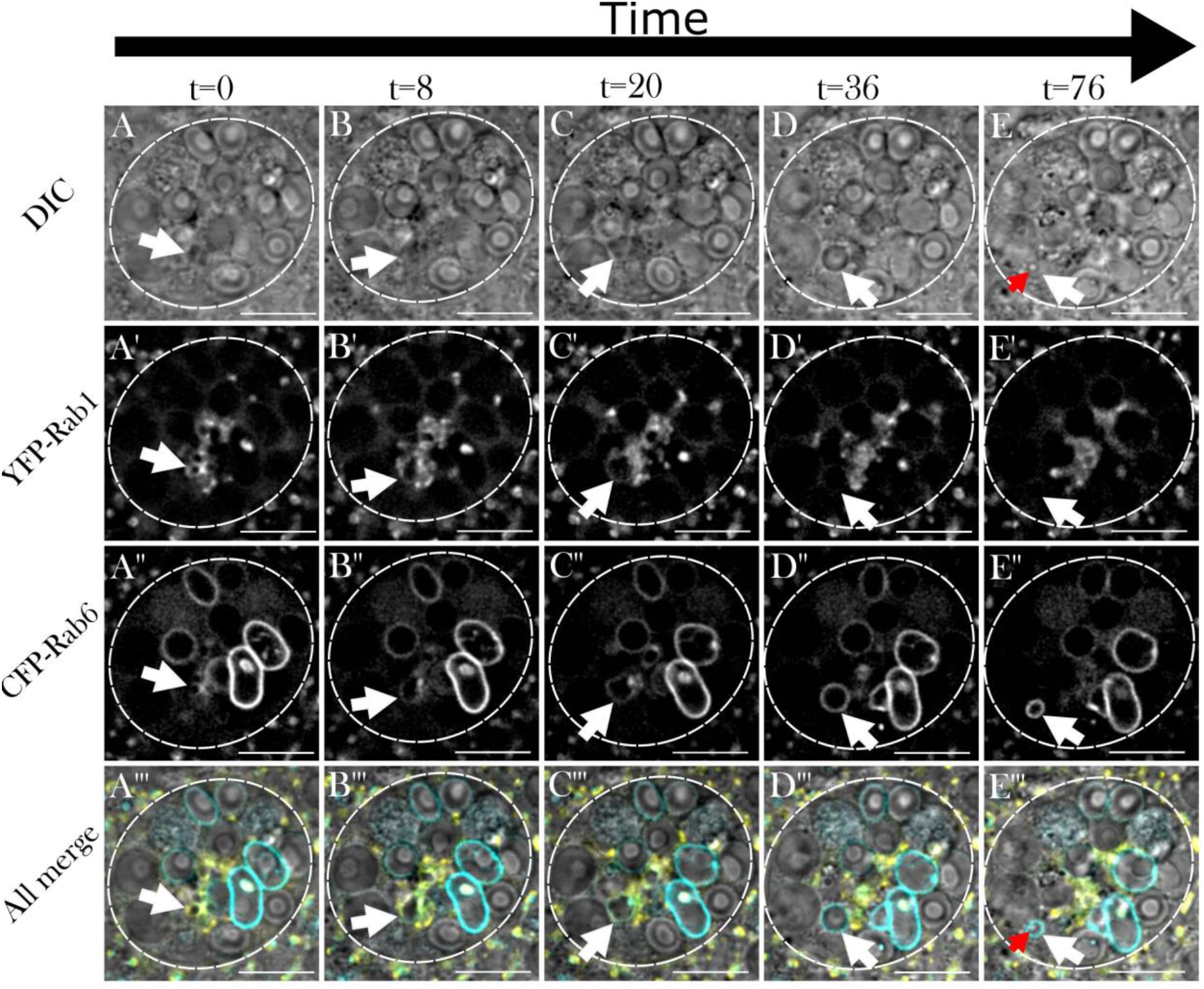
A Rab1 to Rab6 transition accompanies the maturation of secretory compartments at the *trans*-Golgi network of *Drosophila* SCs (related to Movie S1) Panels show *ex vivo* images of a single SC taken at five discrete timepoints with time since start of imaging shown above in minutes. Rows within panel display cellular organisation at each timepoint through DIC imaging (A-E), fluorescent YFP-Rab1 signal (A’-E’), fluorescent CFP-Rab6 signal (A’’-E’’), and combined images displaying all three (A’’’-E’’’). White arrows indicate the position of a secretory compartment as it matures through a Rab1 to Rab6 transition across time and red arrows indicate the position of a newly formed DCG inside that compartment. (A-C) A small, central compartment (white arrow; <1µm diameter) that is primarily labelled with YFP-Rab1 grows rapidly in size, losing most of the YFP-Rab1 signal from its surface and accumulating more CFP-Rab6. (D) The compartment loses all detectable YFP-Rab1 signal from its surface, obtains its greatest diameter, becomes perfectly spherical and starts to migrate peripherally. (E) The compartment retains its CFP-Rab6 identity but begins to contract again in diameter, as a DCG rapidly appears inside it (red arrow). The time interval between the formation of a large Rab6-positive compartment and DCG biogenesis varies between compartments, with this example being particularly rapid. Approximate outline of SC is marked by dashed circles. Scale bars: 10 µm.

We also investigated what mechanisms might permit small Rab1-positive compartments to grow so rapidly. Through further time-lapse imaging, we found that jointly labelled Rab1/Rab6-compartments are able to fuse together, thereby increasing their size as they mature (Supp. Fig. S2, Movie S2). Since later compartments marked solely by Rab6 do not appear to be capable of homotypic fusion, our findings suggest that either Rab1 or a combination of Rab1 and Rab6 are required for these fusion events to occur.

### A Rab6 to Rab11 transition accompanies DCG formation in SCs

We have previously shown that a new DCG compartment is generated every 4-6 h in SCs (Redhai et al., 2016). Having demonstrated that Rab11 marks all DCG compartments and that Rab6 and Rab11 colocalise on some DCG compartments and occasionally on one large non-acidic, core-less compartment, we investigated whether Rab6-positive compartments that lack DCGs might be the precursors of Rab11- marked DCG compartments, using flies expressing both the *CFP-Rab6* and *YFP- Rab11* gene traps.

These experiments showed a clear transition from Rab6- to Rab11-labelling on secretory compartments, a process that occurred over the course of many hours (Fig. 3, Movie S3; Supp. Fig. S3, Movie S4). This transition took place in several stages and coincided with a number of important changes in the compartment. At the earliest stage, compartments marked by Rab6 lack a DCG and have no Rab11 present on their limiting membranes (Fig. 3A, white arrow, and Supp. Fig. S3A). Subsequently, Rab11 begins to be recruited and this coincides with the compartment contracting in size, the beginning of ILV biogenesis, and the appearance of transient dense accumulations of Rab6 on the inside of the limiting membrane (Fig. 3B and Supp. Fig. S3B). It also immediately precedes the formation of DCGs inside compartments (Fig. 3C). As well as occurring soon after the recruitment of Rab11, DCG biogenesis is completed rapidly, typically in less than 20 min (Movie S3). Following DCG biogenesis, maturing secretory compartments continue to recruit Rab11 to their membranes whilst gradually losing Rab6, resulting in DCG compartments that are labelled primarily by Rab11, but may still contain Rab6-positive ILV puncta (Fig. 3D-F and Supp. Fig. S3C), which presumably persists, even after Rab6 has been removed from the compartment’s limiting membrane (Fig. 1I; Supp. Fig. 1D).

**Fig 3.**
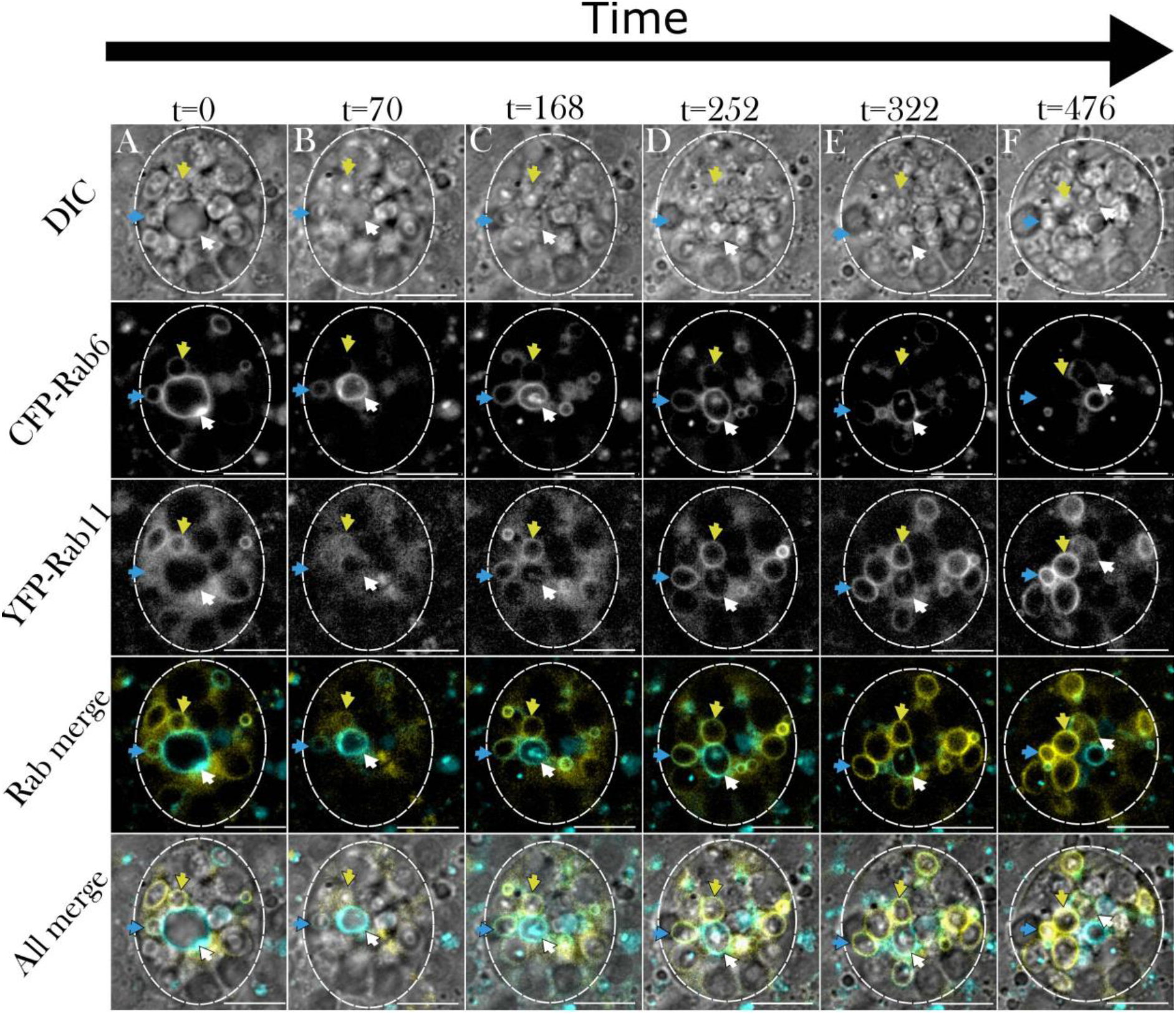
Rab6 to Rab11 transition on surface of maturing secretory compartments in *Drosophila* secondary cells coincides with exosome and DCG biogenesis (related to Movie S3) Panel shows *ex vivo* images of a single SC taken at six discrete timepoints with time since start of imaging shown above in minutes. Rows within panel display cellular organisation at each timepoint through DIC imaging (A-F), fluorescent CFP-Rab6 signal (A’-F’), fluorescent YFP-Rab11 signal (A’’-F’’), a combined fluorescence image (A’’’-F’’’) and a combined DIC and fluorescence image (A’’’’-F’’’’). Three coloured arrows (blue, yellow and white) each indicate the position of one maturing secretory compartment across time. (A-A’’’’) The compartments marked by blue and yellow arrows begin with CFP-Rab6 and YFP-Rab11 co-labelling and have DCGs already present. The compartment marked by the white arrow is significantly larger, is labelled strongly with CFP-Rab6, but has no YFP-Rab11 on its surface and no DCG. (B-B’’’’, C-C’’’’) The blue and yellow arrowed compartments lose CFP-Rab6 labelling over time and become more heavily marked by YFP-Rab11. The compartment marked with a white arrow significantly contracts in size and is only weakly labelled by CFP-Rab6 by the end of the time course. In contrast, YFP-Rab11 begins to accumulate on the compartment. Simultaneously, this compartment begins forming internal structures, with ILVs appearing first (B’) and then a DCG (C). ILVs are marked by both CFP-Rab6 and YFP- Rab11 and at least partly surround the DCG (C). (D-D’’’’, E-E’’’’, F-F’’’’). The two more mature highlighted compartments lose CFP-Rab6 identity and are strongly labelled by YFP-Rab11 by the end of the time course. The compartment labelled with a white arrow still retains some CFP-Rab6, but YFP-Rab11 continues to increase in levels. Approximate outline of SC is marked by dashed circles. Scale bars: 10 µm.

### DCG biogenesis in SCs is regulated by evolutionarily conserved mechanisms

The changes in Rab identity observed when DCG compartments mature in SCs are consistent with some of the Rab identities of secretory compartments at different maturation stages previously reported in other secretory cells in flies and other organisms (Ailion et al., 2014; Cattin-Ortolá et al., 2017; Thomas et al., 2021; Ma and Brill, 2021). Despite the remarkably large size of SC DCGs, we reasoned that DCG compartments in these cells were likely to be assembled via similar mechanisms to those employed in mammalian cells. To test this, we assessed the effects of knocking down two conserved trafficking regulators involved in DCG biogenesis in mammalian cells, *Arf1* (otherwise called *Arf79F*) and components of the AP-1 coatomer complex. Both have established roles in regulating maturation and appropriate cargo loading in DCG compartments (e.g. Stamnes et al., 1993; Traub et al., 1993; Bonnemaison et al., 2013). To mark dense cores, we expressed a GPI-anchored form of GFP, which concentrates in the DCGs of SCs as they mature (Fig. 4A; Redhai et al., 2016).

**Fig 4.**
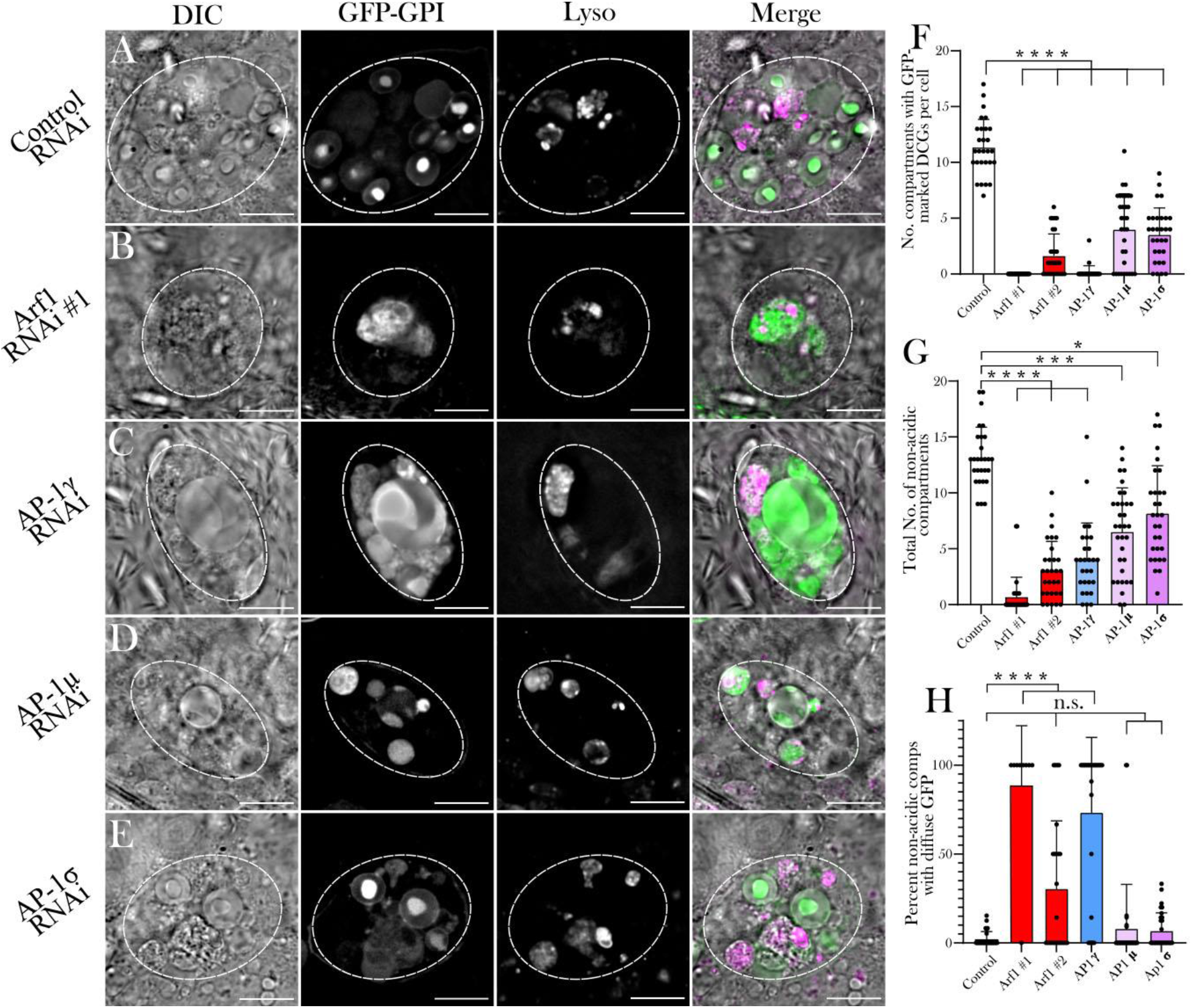
The conserved trafficking regulators Arf1 and AP-1 are essential for DCG biogenesis in SCs (A-E) Representative images of SCs expressing the DCG marker *GFP-GPI* together with a control RNAi (A) or RNAis targeting *Arf1* (RNAi #1; B), *AP-1γ* (C), *AP-1μ* (D) or *AP-1σ* (E). Cellular organisation was assessed using DIC imaging, GFP-GPI fluorescence and Lysotracker Red fluorescence; a merged image is also shown for each cell. Note that the knockdowns generally reduce the number of large non-acidic compartments and the number of DCG compartments, though some remaining compartments can be expanded in size. (F) Histogram showing number of compartments containing GFP-labelled DCGs in control SCs and following knockdown of *Arf1* and *AP-1* subunits. (G) Histogram showing number of non-acidic compartments in these different genotypes. (H) Histogram showing the percentage of non-acidic compartments with diffuse GFP-GPI in these different genotypes. Approximate outlines of SCs are marked by dashed circles. Scale bars: 10 µm. P<0.05: * P<0.01: ** P<0.001: *** P<0.0001: ****.

Knockdown of *Arf1* with two independent RNAis expressed under the control of the *dsx*-GAL4 driver, which in the accessory gland is active only within SCs, almost invariably produced SCs without any DCGs, as judged by both GFP-GPI fluorescence and DIC microscopy (Fig. 4B, F; Supp. Fig. S4B). In addition, significantly fewer non-acidic compartments were produced, which were often larger than normal and which contained diffuse GFP rather than the concentrated cores of GFP fluorescence seen in normal DCGs (Fig. 4B, G, H). Compared to controls, a greater proportion of large acidic late endosomal and lysosomal compartments also contained GFP (Supp. Fig. S5A), presumably reflecting more trafficking of GFP-GPI to these compartments and/or reduced GFP quenching or degradation within them.

Knockdown of three different subunits of the AP-1 coatomer complex, *AP-1γ*, *AP-1µ* and *AP-1σ*, also strongly reduced the number of DCG compartments in SCs and the total number of non-acidic compartments (Fig. 4C-G). In the knockdown producing the most extreme phenotypes, *AP-1γ*, the remaining non-acidic compartments frequently contained diffuse GFP (Fig. 4H). Furthermore, a greater proportion of acidic compartments contained fluorescent GFP (Supp. Fig. S5A).

In mammalian cells, Arf1 and the AP-1 complex are thought to control the formation of DCG compartments and subsequent cargo loading respectively. In view of our findings that this process involves a Rab6 to Rab11 transition in SCs, we tested whether Arf1 and AP-1 control this transition. For this, we knocked down *Arf1* and *AP- 1* components in SCs expressing either the *CFP-Rab6* or *YFP-Rab11* gene traps. Interestingly, although these fly lines drive RNAi expression under the control of *dsx*- GAL4 in the same way as the GFP-GPI line we had used, the level of knockdown appeared to be reduced and some DCG compartments were generated in many of these genetic backgrounds.

Nevertheless, knockdown of *Arf1*, *AP-1γ*, *AP-1µ* and *AP-1σ* in SCs significantly reduced the total number of DCG compartments and the number of Rab11-positive compartments produced (Fig. 5A-G, Supp. Fig. S4D). Interestingly, the percentage of Rab11-compartments which contained DCGs was, for most genotypes, not significantly affected by these knockdowns, with almost all Rab11-compartments still forming DCGs in the majority of SCs (Supp. Fig. S5B). This suggests that DCG biogenesis can still take place in compartments that have undergone the Rab6 to Rab11 transition. Notably however, a greater proportion of the DCGs produced were malformed or irregular, often being split into multiple fragments within a compartment (Fig. 5H), indicating that essential factors for normal DCG biogenesis or maturation were not present at appropriate levels in these compartments due to *Arf1* and *AP-1* knockdowns.

**Fig 5.**
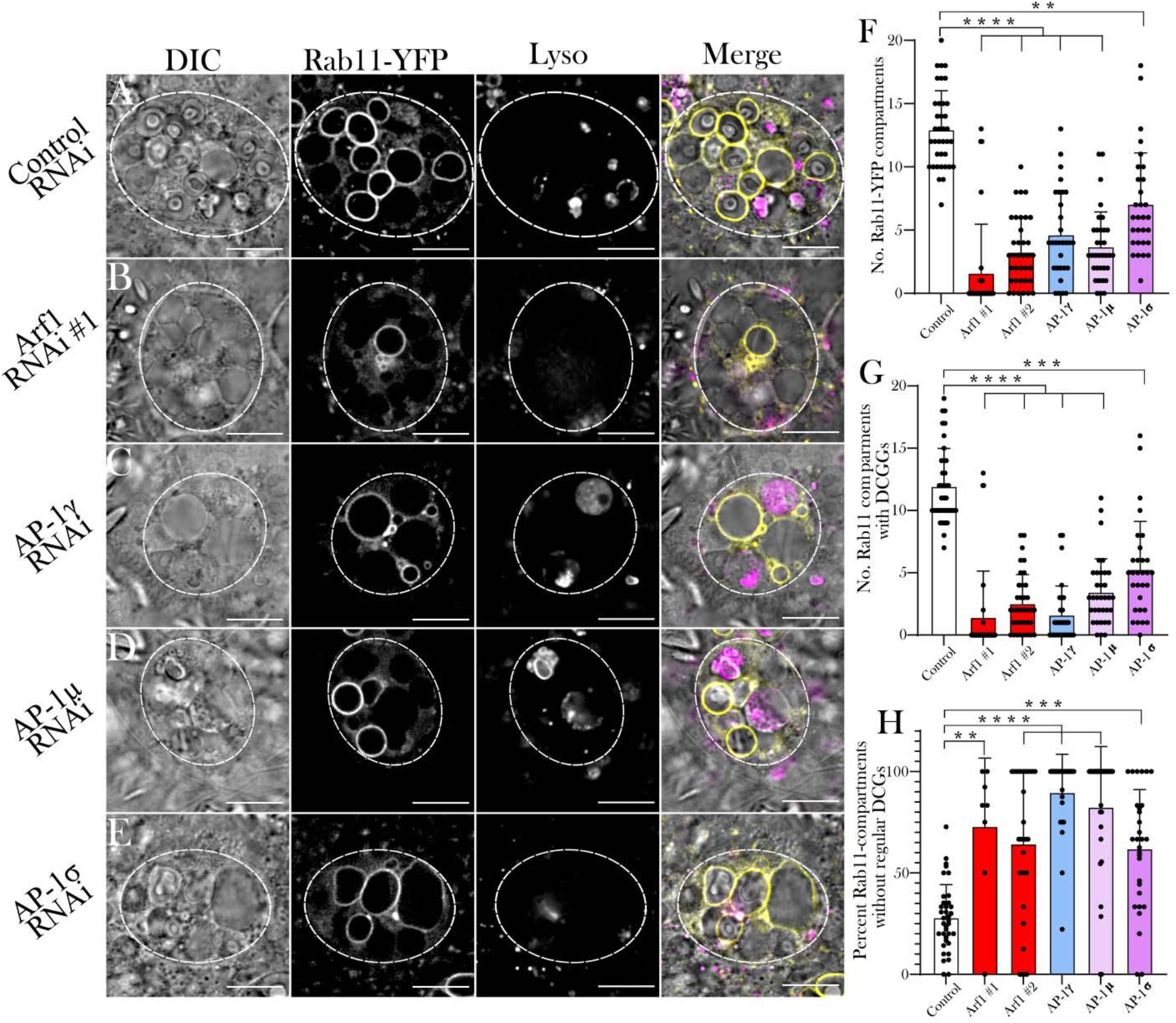
*Arf1* and *AP-1* regulate Rab11-compartment identity and subsequent DCG biogenesis (A-E) Representative images of SCs expressing the *YFP*-*Rab11* gene trap together with a control RNAi (A) or RNAis targeting *Arf1* (RNAi #1; B), *AP-1γ* (C), *AP-1μ* (D) or *AP-1σ* (E). Cellular organisation is assessed using DIC imaging, YFP-Rab11 fluorescence and Lysotracker Red fluorescence, and a merged image is provided for each cell. Note that in the knockdown cells, there are fewer Rab11-positive compartments and more of them either do not contain DCGs, or contain abnormally shaped or multiple DCGs, when compared to controls. (F) Histogram showing number of YFP-Rab11-labelled compartments in control SCs and following knockdown of *Arf1* and *AP-1*. (G) Histogram showing the number of YFP-Rab11 compartments containing DCGs in these different genotypes. (H) Histogram showing the percentage of YFP-Rab11 compartments which fail to produce regularly shaped DCGs in these different genotypes. Approximate outlines of SCs are marked by dashed circles. Scale bars: 10 µm P<0.05: * P<0.01: ** P<0.001: *** P<0.0001: ****

For the two knockdowns that had the most profound effect on DCG number (*Arf1*- RNAi #1 and *AP-1γ-*RNAi; Fig. 5B, C, G), many cells did not contain any DCGs, mirroring the effects of these knockdowns in SCs expressing the GFP-GPI marker (Fig. 4B, C). However, the phenotypes produced by these two knockdowns with the YFP-Rab11 marker differed. While the rare Rab11-positive compartments in *Arf1*- RNAi #1-expressing SCs almost invariably contained DCGs, most Rab11-positive compartments did not in the *AP-1γ* knockdown (Supp. Fig. S5B), consistent with AP- 1 having a key role in maturation of these compartments after the Rab6 to Rab11 transition takes place.

All but one of the different knockdowns also reduced the number of Rab6-positive compartments, particularly the *Arf1*-RNAi #2 knockdown, where only 1.0 ± 1.2 CFP- Rab6-labelled compartments were observed (Fig. 6A-E, H, Supp. Fig. S4H). Interestingly, in both *Arf1* knockdowns, but not in *AP-1* subunit knockdowns, a significant proportion of large non-acidic, core-less compartments were not strongly labelled with CFP-Rab6 (Supp. Fig. S5D), suggesting that they had not matured to Rab6 identity. Indeed, when *Arf1* was knocked down in SCs expressing the *YFP-Rab1* gene trap, a number of these large non-acidic, non-DCG compartments were marked by Rab1, unlike in controls (compare Fig. 6G and 6F), indicating that these compartments were unable to transition to a Rab6-positive identity and instead remained as Rab1-positive compartments. The existence of these large compartments marked by Rab1 also suggests that Rab6 is not required for the fusion events between compartments formed in the *trans*-Golgi and compartment expansion to occur. Interestingly, despite having a related distribution to YFP-Rab1 in control cells, we found no evidence of YFP-Rab2 acting as a marker on these enlarged compartments produced by *Arf1* knockdown (Supp. Fig. S4H)

**Fig 6.**
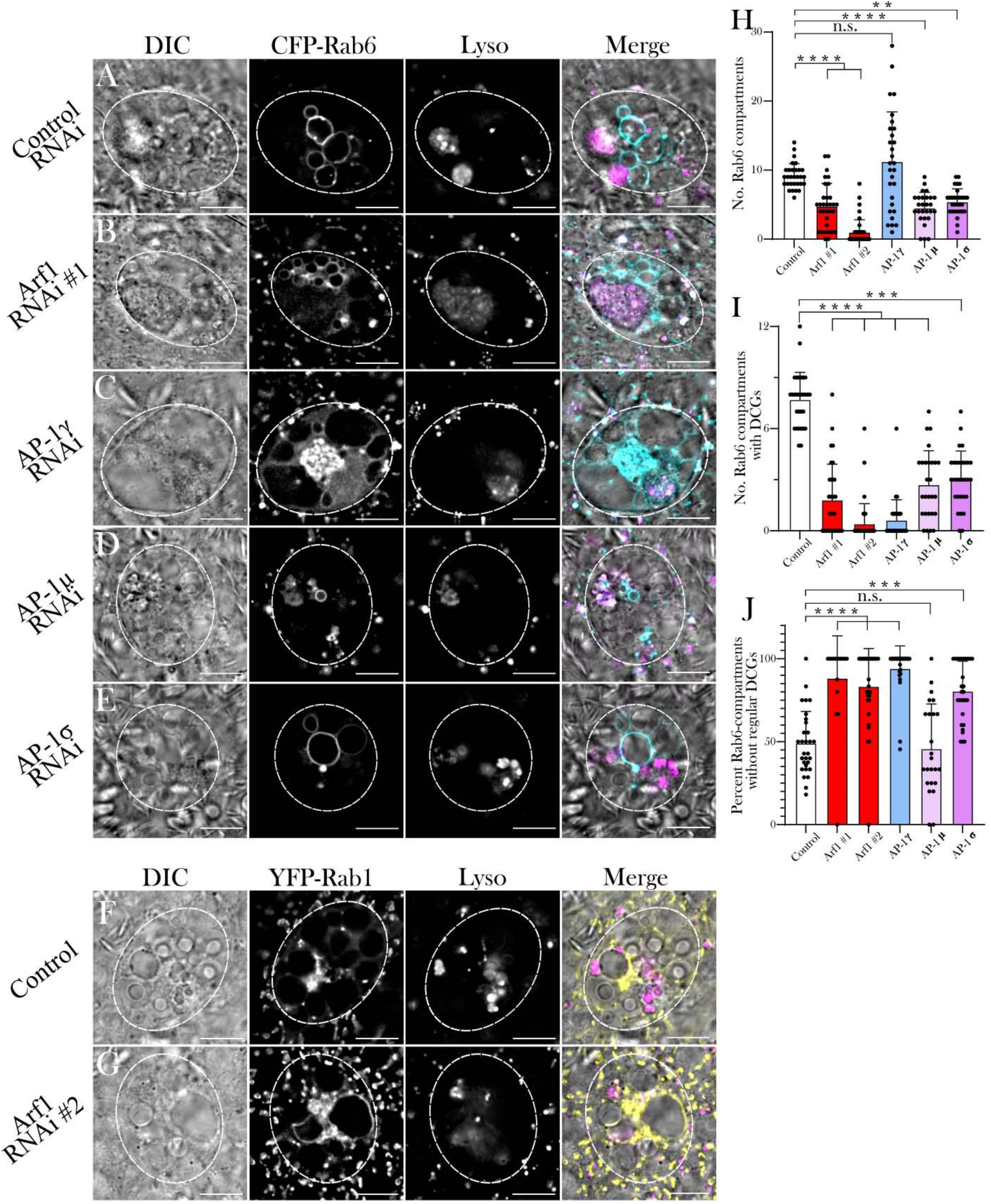
*Arf1* and *AP-1* regulate Rab6-compartment identity and the maturation of DCG compartments (A-E) Representative images of SCs expressing the *CFP*-*Rab6* gene trap together with a control RNAi (A) or RNAis targeting *Arf1* (RNAi #1; B), *AP-1γ* (C), *AP-1μ* (D) or *AP-1σ* (E). Cellular organisation is assessed through DIC imaging, CFP-Rab6 fluorescence and Lysotracker Red fluorescence, and a merged image is presented for each cell. Note the number of CFP-Rab6-positive compartments is reduced in all knockdown backgrounds, except *AP-1γ*, and in all knockdowns, few labelled compartments contain DCGs. (F, G) Representative SCs co-expressing the *YFP- Rab1* gene trap either alone (F) or together with an *Arf1* RNAi #2 (G). Note that in control *YFP-Rab1* SCs, no large non-acidic compartments are Rab1-positive. When SCs are subjected to *Arf1* knockdown, by contrast, YFP-Rab1 does mark several large non-acidic compartments which do not contain DCGs. (H) Histogram showing number of CFP-Rab6-labelled compartments in control SCs and following knockdown of *Arf1* and *AP-1*. (I) Histogram showing number of CFP-Rab6-compartments containing DCGs in these different genotypes. (J) Histogram showing the percentage of CFP- Rab6-compartments, which fail to produce regularly shaped DCGs in these different genotypes. Approximate outlines of SCs are marked by dashed circles. Scale bars: 10 µm. P<0.05: * P<0.01: ** P<0.001: *** P<0.0001: ****.

Consistent with Arf1 and AP-1 being directly or indirectly involved in the Rab6 to Rab11 transition during DCG compartment maturation, a smaller number and proportion of Rab6-positive compartments contained DCGs for all knockdowns compared to controls (Fig. 6I; Supp. Fig. 5C), suggesting that *Arf1* and *AP-1* knockdowns prevent the Rab6 to Rab11 transition that accompanies DCG biogenesis. Likewise, as observed in the YFP-Rab11 background, a significantly higher proportion of the DCGs that were produced were irregular and/or fragmented in most knockdowns (Fig. 6J). The *AP-1γ* knockdown, in which, unlike other knockdowns, the average number of Rab6-positive compartments was not reduced compared to controls (Fig. 6H), was particularly notable. The number of Rab11-positive compartments was strongly reduced (Fig. 5F), and, in most cells, none of the compartments contained DCGs (Fig. 6I). This is consistent with AP-1 playing a key role in the Rab6 to Rab11 transition, and the hypothesis that this transition is required for DCG biogenesis.

In summary, homologues of Arf1 and the AP-1 subunits play a critical role in DCG biogenesis in SCs, as has also been reported in mammalian cells. Our data suggest that Arf1 acts earlier in the process than AP-1, since unlike AP-1, it is required to recruit Rab6 to large compartments, as well as for the later conversion to Rab11-positive compartments. By contrast, AP-1 is primarily involved in the Rab6 to Rab11 transition and in subsequent maturation events that induce DCG formation. The wide range of genetic and imaging tools available to analyse the large secretory compartments of SCs has allowed us to show that their importance in some of the processes during maturation of DCG progenitor compartments and formation of DCGs differs.

### The Rab6 to Rab11 transition is required for DCG and exosome biogenesis

In light of the correlation between DCG biogenesis and the Rab6 to Rab11 transition, we hypothesised that this transition might be required for the assembly of DCGs. To test this, two independent RNAis were used to knockdown each of *Rab6* and *Rab11* in SCs. This allowed us to determine for what processes in DCG biogenesis each Rab was required. Where both Rabs were found to be necessary for a process to occur, we concluded that the Rab6 to Rab11 transition was required to facilitate that process.

Inducing knockdown of either *Rab6* or *Rab11* in SCs expressing the GFP-GPI DCG marker leads to a strong reduction in the number of DCGs present in SCs (Fig. 7A-D; Supp. Fig. S6A-C). Alongside this reduction in DCG-containing compartments, significantly fewer non-acidic compartments were present and many of the remaining non-acidic compartments were marked by diffuse GFP, just as we observed following *Arf1* and *AP-1* subunit knockdown in the GFP-GPI background (Fig. 7E, F). Likewise, the proportion of acidic compartments containing fluorescent GFP increased significantly following either *Rab6* or *Rab11* knockdowns (Supp. Fig. S6D), suggesting that these knockdowns either increase trafficking of secretory compartments to the lysosomal pathway or disrupt lysosome activity. These results suggested that both Rab6 and Rab11 are required for DCG biogenesis.

**Fig 7.**
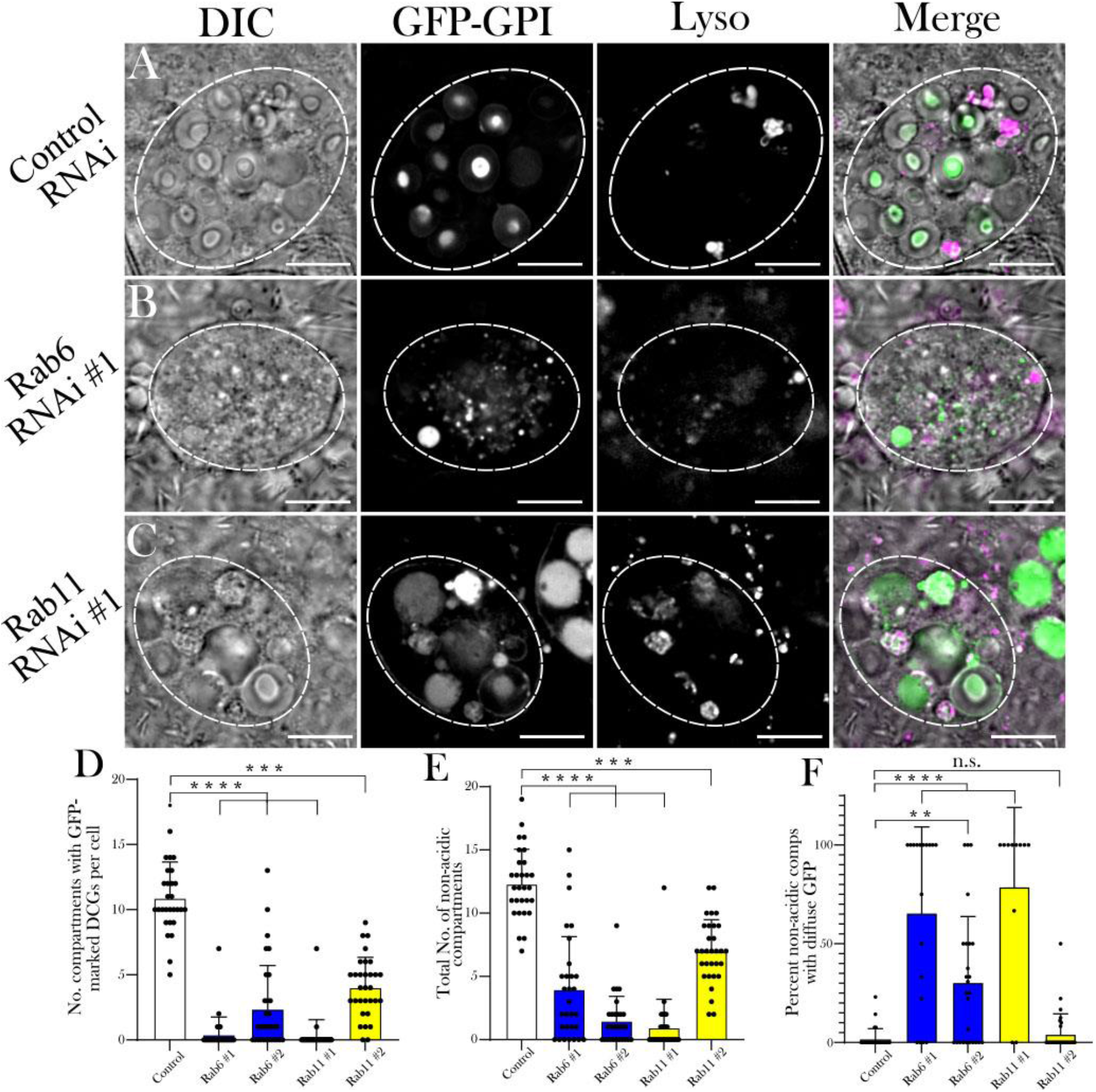
Rab6 and Rab11 are both required for DCG biogenesis in SCs (A-C) Representative images of SCs expressing the DCG marker *GFP-GPI* with a control RNAi (A) or RNAis targeting *Rab6* (B) or *Rab11* (C). Cellular organisation is assessed by DIC imaging, GFP-GPI fluorescence and Lysotracker Red fluorescence, and a merged image for each cell. Note the reduction in large non-acidic compartments in these backgrounds with fewer containing DCGs and in some cases, the remaining compartments often filled with diffuse GFP. (D) Histogram showing the number of compartments containing GFP-labelled DCGs in control SCs and following knockdown of *Rab6* and *Rab11*. (E) Histogram showing the number of large non-acidic compartments in these different genotypes. (F) Histogram showing the percentage of large non-acidic compartments with diffuse GFP-GPI present in these different genotypes. Approximate outlines of SCs are marked by dashed circles. Scale bars: 10 µm. P<0.05: * P<0.01: ** P<0.001: *** P<0.0001: ****.

To better understand these findings, we expressed the same *Rab6* and *Rab11* RNAis in the *CFP-Rab6* and *YFP-Rab11* backgrounds. Just as seen in the GFP-GPI background, all knockdowns significantly reduced the number of Rab6- and Rab11- marked compartments forming DCGs, except for knockdown with *Rab11-*RNAi #2, which increased the total number of Rab6-compartments, leading to the number of DCG-containing Rab6-compartments being unaffected (Fig. 8 A-H, L; Supp. Fig. S7). Interestingly, the loss of DCGs in the *Rab* gene trap backgrounds was not as severe as the loss observed in the GFP-GPI background. This mirrored our findings with *Arf1* and *AP-1* subunit knockdowns, where we concluded that knockdown of these molecules was reduced in these Rab marker lines. This conclusion is supported by the fact that a small number of compartments marked by YFP-Rab11 and CFP-Rab6 were still present even after knockdown of the corresponding *Rab* (eg, Fig. 8C, E, Supp. Fig. 7C) with *Rab11*-RNAi #2 and *Rab6*-RNAi #1 being less effective than the other RNAi used for these two *Rabs*.

**Figure 8.**
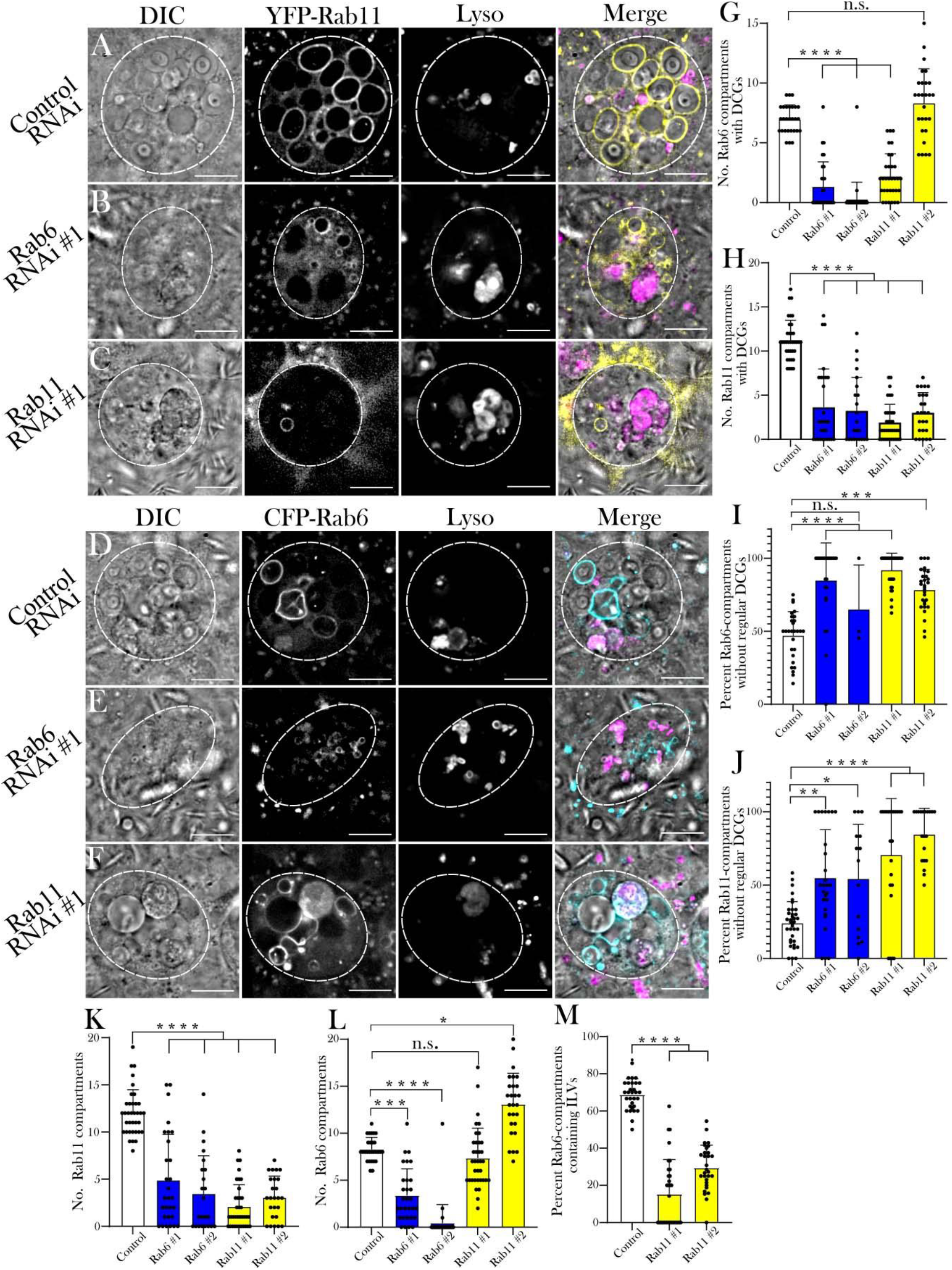
The Rab6 to Rab11 transition on secretory compartments controls exosome as well as DCG biogenesis in SCs (A-C) Representative images of SCs expressing the *YFP*-*Rab11* gene trap with a control RNAi (A) or RNAis targeting *Rab6* (B) or *Rab11* (C). (D-F) Representative images of SCs expressing the *CFP*-*Rab6* gene trap with a control RNAi (D) or RNAis targeting *Rab6* (E) or *Rab11* (F). Cellular organisation in all genotypes is assessed through DIC imaging, gene trap fluorescence and Lysotracker Red fluorescence, as well as a merged image for each cell. Note that when *Rab6* is knocked down, there are reduced numbers of Rab11-compartments and fewer of those that remain contain normal DCGs (H, J, K). By contrast, *Rab11* knockdown does not reduce the number of Rab6-positive compartments, but fewer of these compartments contain normal DCGs or ILVs (G, I, L, M). Also, note that some Rab11 gene trap fluorescence is still visible even after knockdown of *Rab11* (C), and similarly for CFP-Rab6 in the *Rab6* knockdown (E). (G) Histogram showing the number of CFP-Rab6-compartments containing DCGs in control SCs and following *Rab6* and *Rab11* knockdown. (H) Histogram showing the number of YFP-Rab11-compartments containing DCGs in these different genotypes. (I) Histogram showing the percentage of CFP-Rab6 compartments which fail to produce regular DCGs in these different genotypes. (J) Histogram showing the percentage of YFP-Rab11 compartments which fail to produce regular DCGs in these different genotypes. (K) Histogram showing the number of YFP- Rab11-compartments in these different genotypes. (L) Histogram showing the number of CFP-Rab6 compartments in these different genotypes. (M) Histogram showing the percentage of CFP-Rab6 compartments which contain CFP-Rab6-labelled ILVs following knockdown of *Rab11* in SCs versus controls. Approximate outlines of SCs are marked by dashed circles. Scale bars: 10 µm. P<0.05: * P<0.01: ** P<0.001: *** P<0.0001: ****.

We next investigated whether those DCGs that did form after *Rab* knockdown matured normally, by analysing the number of compartments that contained either immature/irregular DCGs or no DCG at all. We found that knockdown of *Rab6* or *Rab11* greatly increased the proportion of compartments which lack a mature, regular DCG, both in CFP-Rab6- and YFP-Rab11-labelled compartments (Fig. 8 I, J), with many cells not containing any normal DCGs. This included knockdown with leaky *Rab11*-RNAi #2 (Fig. 8G), where approximately 75% of Rab6-compartments were either core-less or contained irregular DCGs (Fig. 8I). Overall, these results mirror those for *Arf1* and *AP-1* knockdowns and indicate that both Rab6 and Rab11 are required for DCG biogenesis, since knockdown of either *Rab* reduces the total number of DCGs (Fig. 7D, Fig. 8G, H) and decreases the proportion of large secretory compartments that can form a mature regularly shaped DCG (Fig. 8I, J).

In order to further assess whether the compartment transition from Rab6- to Rab11- positive identity is required for DCG biogenesis, we also looked at the effect of *Rab6* and *Rab11* knockdown on secretory compartment identity. Following knockdown of *Rab6* in a *YFP-Rab11* background, we observed a very large reduction in the number of Rab11-positive compartments, with an approximately 75% decline in these compartments relative to controls and many cases where no Rab11-compartments were formed at all in SCs (Fig. 8K). Importantly, this was accompanied by the large decline in mature DCGs discussed earlier (Fig. 8H, J). In contrast, knockdown of *Rab11* in a *CFP-Rab6* background also prevented normal DCG formation, but had no discernible effect on the number of Rab6-positive compartments or even increased their number (Fig. 8L). These results are consistent with our finding that a Rab6 to Rab11 transition occurs as secretory compartments mature, and that this is required for the production of DCGs.

Finally, having shown that Rab11 is required for DCG formation, we examined its role in biogenesis of exosomes produced in Rab11-compartments, collectively termed Rab11-exosomes. As shown earlier, whilst imaging control samples in the *CFP-Rab6* background, we had noted the very strong correlation between the presence of DCGs and Rab6-positive ILVs within compartments, with these ILVs almost exclusively appearing in DCG compartments (Fig. 1M), from where they can be secreted as Rab11-exosomes. Since time-lapse imaging had shown that ILVs and DCGs appear in compartments within minutes of each other, we investigated the hypothesis that the Rab6 to Rab11 transition is also the trigger for ILV (and consequently Rab11- exosome) biogenesis. Following knockdown of *Rab11* in a *Rab6-CFP* background, a significantly smaller proportion of Rab6-positive compartments produced Rab6- labelled ILVs than in controls (Fig. 8M). As with DCG formation, this result indicates that the recruitment of Rab11 to the membrane of compartments derived from the *trans*-Golgi network is an important step for biogenesis of Rab11-exosomes.

## Discussion

The genetic dissection of DCG biogenesis in the regulated secretory pathway has been restricted by availability of *in vivo* models and the limited possibilities to employ fluorescence and real-time imaging to study the processes involved. Here, we employ SCs of the *Drosophila* male accessory gland to overcome these hurdles. We demonstrate that some of the best characterised regulators of DCG biogenesis in mammals are also involved in DCG formation in SCs and show that a series of Rab transitions involving Rab1, Rab6 and Rab11 precede DCG formation. These transitions suggest a critical interaction between the secretory and recycling endosomal pathways, which is controlled by trafficking regulators like Arf1 and AP-1 (see model in Fig. 9). Our studies provide a platform for more extensive genetic analysis of DCG biogenesis, which should inform our understanding of this process in human health and disease.

**Figure 9.**
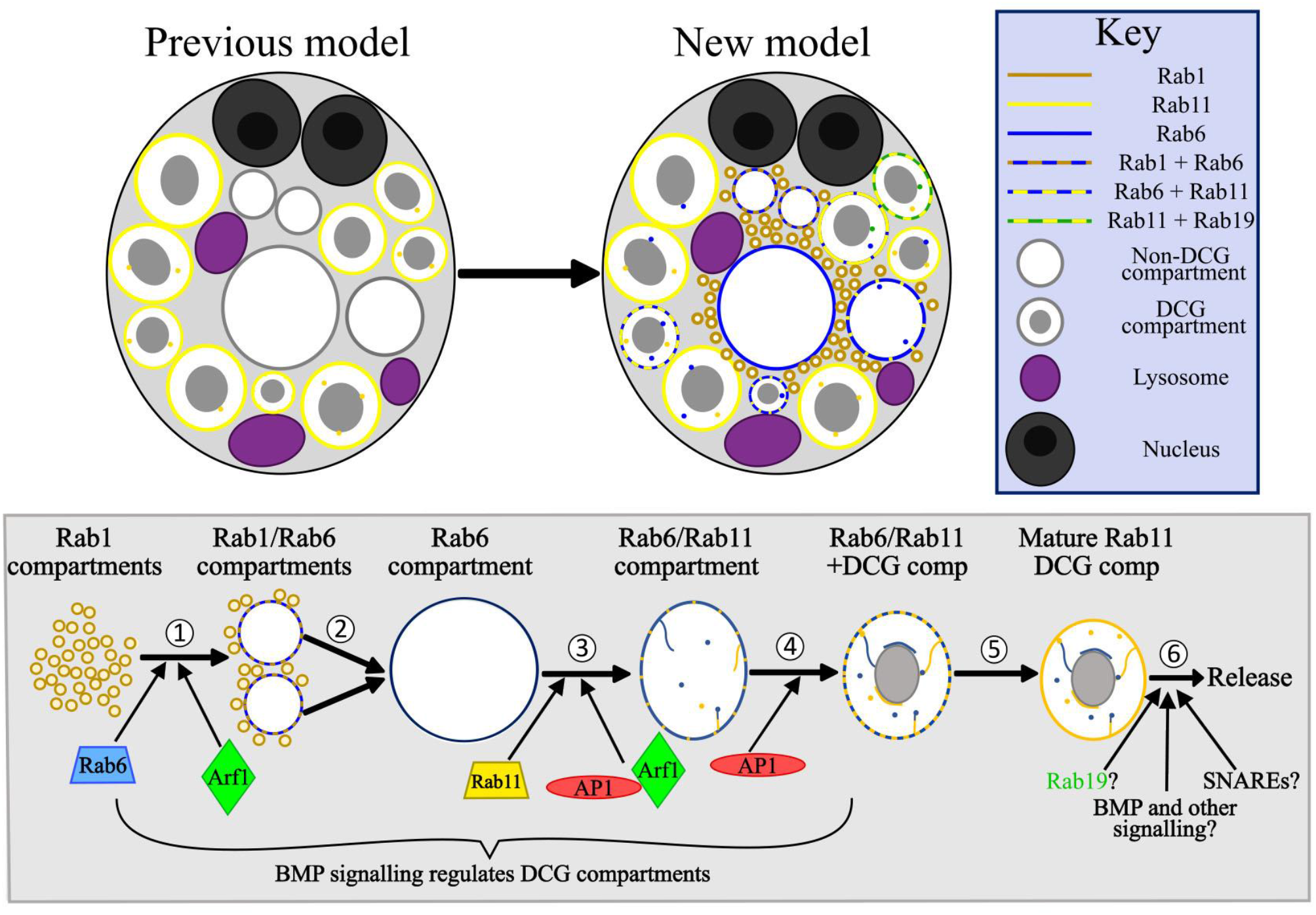
Model for the regulation of DCG compartment biogenesis in SCs (A) A schematic illustrating our previous and updated model of SC secretory and endosomal compartment organisation. Whereas previously it was recognised that DCG compartments in SCs were labelled by Rab11, we have now shown that Rab6 and Rab19 also mark large secretory compartments and can co-label compartments alongside Rab11. We have also demonstrated that Rab1 marks a population of smaller compartments near the cell centre and can colocalise with Rab6 on the surface of growing compartments. Finally, as well as the Rab11-positive ILVs which were recognised beforehand, we have also described the existence of ILVs marked by Rab6 and Rab19 which can be found in compartments labelled by Rab11, and will be secreted as Rab11-exosomes. (B) Schematic outlining the genetic regulation of secretory compartment maturation and DCG biogenesis in SCs. Our results have highlighted at least 6 discrete stages which occur during secretory compartment maturation. (1) In the earliest stage, small Rab1-compartments fuse together and recruit Rab6 to their limiting membrane, creating enlarged Rab1/Rab6-positive compartments. (2) These compartments continue to grow, at least in part via fusion events, until they eventually lose all Rab1 identity and are marked solely by Rab6, which contain no DCGs and no ILVs. The Rab1 to Rab6 transition is regulated by Arf1 and Rab6 recruitment is required to progress to later maturation steps. (3) Soon after their formation, Rab6-positive compartments contract in size. They subsequently recruit Rab11 to their limiting membrane which induces the formation of ILVs, some of which appear to coalesce into the long ILV chains we have observed (internal lines in compartments). The recruitment of Rab11 is regulated by Arf1 and the AP-1 complex, without which most Rab11-compartments fail to form. Any that do form usually do not mature normally. (4) As Rab11 continues to be recruited to membranes, DCG biogenesis occurs within compartments. Redhai et al. (2016) previously demonstrated that BMP signalling regulates the rate of DCG- compartment biogenesis, indicating that BMP acts at one or more points upstream of this event. We additionally show here that AP-1 regulates normal DCG biogenesis. (5) Over several more hours, Rab6 is fully shed from the limiting membrane, leaving a matured secretory compartment which contains a DCG and a mix of ILVs. (6) Matured compartments are eventually secreted following fusion with the plasma membrane. Secretory compartment release was previously shown to be regulated by BMP signalling (Redhai et al., 2016), but what other factors are involved remains unclear. Specific SNARE proteins are likely to be required for compartment fusion to the plasma membrane, and compartments may undergo further maturation steps, possibly regulated by factors such as Rab19 and additional signalling pathways.

### Rab1 and Rab6 mark secretory compartments prior to DCG formation

By expressing fluorescent *Rab* gene-traps in SCs we were able to produce detailed time-lapse videos of maturing secretory compartments. These showed that both Rab1 and Rab6 are present on compartments produced by the *trans*-face of the Golgi at successive stages of maturation, with small Rab1-positive compartments transitioning into larger Rab6-positive compartments over the course of approximately 40 minutes (Figs. 2, 9 and Supp. Fig. S2). After the Rab1-Rab6 transition, when Rab1-YFP signal is no longer visible on the limiting membrane of compartments, a DCG can then form (Fig. 2). This conclusively demonstrates that these Rab1/Rab6-marked compartments are the same structures that go on to form DCGs. Our results fit well with previous findings that Rab1 drives secretory granule maturation in *Drosophila* salivary glands and with observations that Rab1 and Rab6 are located at the periphery of immature granule-forming compartments (Ma & Brill, 2021; Neuman et al., 2021), although in salivary glands, Rab1 may remain associated with more mature compartments. Our time-lapse videos also provided insights into the mechanisms of secretory compartment maturation, showing that multiple smaller Rab1-/Rab6-labelled compartments can fuse to create the larger Rab6-compartments in SCs.

### The Rab6 to Rab11 transition accompanies DCG biogenesis and is modulated by known DCG regulators

Through further time-lapse imaging, we found that another Rab transition takes place on the surface of secretory compartments prior to DCG biogenesis. Rab6 on the surface of compartments is gradually replaced by Rab11 over the course of many hours, with DCG biogenesis occurring minutes after the beginning of this transition (eg. Fig. 3). From the time-lapse videos it appeared that the entire process of DCG formation was typically complete within 20 minutes.

We also showed, through SC-specific knockdown of *Arf1* and each of the *AP-1* subunits, that the roles of both Arf1 and AP-1 in DCG biogenesis are conserved in SCs. Both Arf1 and AP-1 are evolutionarily conserved trafficking regulators that are known to be required for maturation of DCG compartments (Stamnes et al., 1993; Traub et al., 1993; Bonnemaison et al., 2013). *Arf1* and *AP-1* knockdowns in SCs expressing a range of markers led to significantly reduced DCG formation (Fig. 4). Additionally, experiments in *CFP-Rab6* and *YFP-Rab11* backgrounds showed that knockdown of *Arf1* and *AP-1* subunits significantly decreased the number of both Rab6-positive and Rab11-positive compartments in almost all cases (Figs. 5, 6). Of those compartments that remained, a significantly larger proportion failed to form regular DCGs.

Based on our analysis, we suggest that Arf1 and AP-1 contribute to the process of DCG biogenesis at three different stages. Firstly, Arf1 seems to be required at the Rab1 to Rab6 transition stage, thereby explaining the presence of Rab1-positive, Rab6-negative, large non-DCG compartments in *Arf1* knockdowns. Secondly, since Rab11-compartments and DCG biogenesis are also strongly reduced in *Arf1* and *AP- 1* knockdowns, we propose that their protein products are required during the Rab6 to Rab11 transition and associated DCG formation. Thirdly, even if this transition takes place and DCGs form, likely because knockdown does not completely eliminate all target gene expression, *AP-1* knockdown in particular seems to be required for the establishment of regular DCG morphology (eg. *AP-1γ* knockdown; Fig. 6H-J). One explanation for these latter two roles in DCG biogenesis is that Arf1 and AP-1 direct the fusion and release of vesicles to and from Rab6-/Rab11-positive compartments, thereby delivering materials, which are needed for DCG formation, and removing those that are not (Fig. 9).

### The Rab6 to Rab11 transition is required for DCG biogenesis

Knockdown of either *Rab6* or *Rab11* effectively eliminated DCG formation in SCs, demonstrating that both Rabs are required for granule biogenesis (Fig. 7). Of these, it was particularly significant that *Rab11* knockdown inhibited DCG formation, since this was consistent with our observations that the start of DCG biogenesis occurs after the beginning of the Rab6 to Rab11 transition. The critical role for *Rab11* at this transition stage was supported by the observation that the numbers of Rab6-positive precursor compartments were not reduced by *Rab11* knockdown (Fig. 8L). Together with the observation that DCG biogenesis occurs early in the Rab6 to Rab11 transition, these results strongly suggest that the switch to Rab11 identity acts as the regulatory trigger for DCG biogenesis in SCs.

Rab11 is frequently associated with recycling endosomes (Stenmark, 2009; van IJzendoorn, 2006). However, it is also associated with secretory granules, such as Epidermal Lamellar Granules and the secretory granules of *Drosophila* salivary glands (Ishida-Yamamoto et al., 2007; Ma & Brill, 2021). Likewise, Rab11 is also known to contribute to insulin granule exocytosis in pancreatic β cells (Sugawara et al., 2009), as well as to the trafficking and release of secretory granules in the fungus, *Aspergillus nidulans* (Pinar and Peñalva, 2021). It therefore seems likely that in these other cells, the association with Rab11 has a role to play in the proper maturation of the secretory compartments they produce and in DCG biogenesis, as we have found in SCs.

One important unanswered question is whether the Rab6 to Rab11 transition is associated with delivery of cargos specifically from Rab11-positive recycling endosomes that fuse with the large Rab6-compartments and promote DCG biogenesis. Our data suggest that the GFP-GPI marker enters DCG precursor compartments before the Rab6 to Rab11 transition, because knockdowns that block this transition lead to accumulation of diffuse GFP-GPI in the resulting non-acidic compartments that are formed. However, recycling endosomes are known to be more acidic than the secretory compartments that form at the surface of the *trans*-Golgi.

Therefore, one possibility is that the recycling endosomal system delivers the V-ATPase proton pump or more acidic luminal contents during the Rab6 to Rab11 transition, which might lead to the pH changes in secretory compartments that are known to drive DCG assembly (Wu et al., 2001).

### Rab11-exosome and DCG biogenesis may be interdependent processes controlled by the Rab6 to Rab11 transition

Because of the unique biology of the SC system, we were able to examine additional aspects of DCG compartment regulation that we would not easily be able to investigate in other available models. One key feature was the association of ILVs with the developing DCG inside the maturing compartment. Imaging of control cells in a *CFP- Rab6* background showed that ILVs are almost exclusively found in compartments with DCGs (Fig. 1M). Furthermore, consistent with previous observations (Fan et al., 2020), time-lapse imaging and single wide-field fluorescence micrographs demonstrated that ILVs marked by Rab19-YFP, Rab11-YFP and Rab6-CFP were closely associated with DCGs. Rab11 is an established marker of exosomes secreted from compartments marked by this recycling endosomal Rab, but both Rab19 (Gonzales et al., 2009; Reinhardt et al., 2013) and Rab6 (Buschow et al., 2010; Lazar et al., 2015) have also been reported to be secreted in mammalian extracellular vesicles, suggesting that exosomes labelled by these Rabs are not uniquely produced by SCs.

Extended lengths of many DCG boundaries were clearly marked by Rab-labelled ILVs and the fluorescent Rab-signal revealed long chains of ILVs running from the limiting membranes of compartments to DCGs and then along DCG boundaries. Indeed, it was often possible to discern the position and shape of DCGs through just the distribution of fluorescent ILVs. In addition, we also found that knockdown of *Rab11* significantly reduced the formation of Rab6-positive ILVs, which normally contribute to the population of Rab11-exosomes originating from SC Rab11-compartments, thereby demonstrating that Rab11 is required for the biogenesis of these ILVs as well as DCGs (Fig. 9). These results indicate that ILV formation and DCG biogenesis share common regulatory mechanisms.

A previous study has suggested that knockdown of some components of the core ESCRT complexes disrupt DCG biogenesis in SCs (Marie et al., 2023). Importantly, however, not all *ESCRT* knockdowns have the same effect, suggesting that the formation of ILVs itself may not be the essential event providing this link. Previous studies in SCs have shown that vesicle-associated GAPDH activity regulates changes in ILV clustering which are closely linked with DCG morphology (Dar et al., 2021). Together with these earlier findings, our results highlight the link between ILVs and DCGs and emphasise the need to understand the role of GAPDH and other membrane-associated molecules in regulating this interaction. Furthermore, they suggest that the conserved trafficking pathway that makes Rab11-exosomes (Fan et al., 2020; Marie et al., 2023) may also be critical in producing DCGs in secretory cells.

Finally, both DCG biogenesis and secretion in SCs is regulated by autocrine BMP signalling mediated by the BMP ligand, Decapentaplegic, which is packaged into the DCGs, providing a mechanism for accelerating DCG production when secretion rates are high (Redhai et al., 2016). It will now be interesting to determine which maturation events in SC DCG biogenesis are controlled by BMP signalling (Fig. 9), and whether additional intracellular signalling cascades can modulate this process in other ways. Indeed, given the parallels in the regulation of DCG biogenesis between mammalian cells and fly SCs, it seems likely that as this process is genetically dissected in SCs, it will further inform our understanding of DCG biogenesis mechanisms more generally, and provide insights into human pathologies that involve DCGs.

## Materials and Methods

### Fly stocks

UAS-RNAi lines were sourced from the Bloomington *Drosophila* Stock Centre TRiP collection (BDSC; Ni et al., 2009) and Vienna Drosophila Resource Centre shRNA, GD and KK libraries (VDRC; Dietzl et al., 2007): *rosy-RNAi* as a control (BDSC; 44106; HMS02827; Marie et al., 2020); *Arf1-RNAi* #1 (VDRC; 23082; Wang et al., 2017) and #2 (VDRC; 103572; Wang et al., 2017); *AP-1γ-RNAi* (BDSC; 27533; JF02684; Peterson and Krasnow, 2015); *AP-1µ-RNAi* (VDRC; 24017; Bellec et al., 2021); *AP-1σ-RNAi* (VDRC; 107322); *Rab6-RNAi* #1 (BDSC; 27490; JF02640; Ayala et al., 2018) and #2 (BDSC; 35744; HMS01486); *Rab11 RNAi* #1 (BDSC; 27730; JF02812; Ma and Brill, 2021) and #2 (VDRC; 108382; Nie et al., 2019). We additionally used the following fluorescent gene trap lines: *YFP-Rab11*, *YFP-Rab1*, *YFP-Rab19* (Dunst et al., 2015) and *CFP-Rab6*, provided by S. Eaton and F. Karch; *UAS-GFP-GPI* (Greco et al., 2001; Redhai et al., 2016). The *CFP-Rab6* and *YFP- Rab11* gene traps and *UAS-GFP-GPI* lines were combined with *tub-GAL80^ts^* (BDSC 7108) and *dsx-GAL4* (provided by S. Goodwin) to produce *Drosophila* lines with one of these three fluorescent markers, as well as the GAL4-GAL80^ts^ machinery that allows SC-specific temperature-inducible expression of UAS-transgenes.

### Fly culture and handling

Flies were cultured on standard cornmeal agar food, using a 12 hour light/dark cycle. Flies carrying UAS-transgenes were crossed with flies which contained the dsx- Gal4/tub-Gal80^ts^ driver system as well as one of the fluorescent gene traps or *UAS-GFP-GPI* and kept at 25°C. Virgin male offspring from this cross were collected on the same day they eclosed and were transferred to 29°C for 6 days to trigger transgene expression. In experiments where no UAS-transgene was expressed and only gene traps were employed, the same timings and temperatures were used, except for time-lapse experiments where incubation at 29°C could vary from 5-7 days post-eclosion.

### Imaging, deconvolution and time-lapse movies

To visualise SC organisation, accessory glands were dissected into cold PBS, incubated with 500nM Lysotracker Red DN-99 (Invitrogen, L7528) for 5 minutes on ice and washed again with cold PBS. Finally, glands were mounted between two coverslips (thickness No. 1.5H, Marienfeld-Superior) in a small drop of PBS. Cover slips were placed into a custom-made metal mount for support. Excess PBS was drawn off using filter paper until glands were slightly flattened between the two coverslips.

Live SCs were imaged using the DeltaVision Elite system from Olympus AppliedPrecision, an inverted wide-field fluorescence microscope that can perform both fluorescence and differential interference contrast (DIC) microscopy. Cells were viewed at 1000X magnification with three SCs imaged and analysed from each accessory gland. SCs were imaged using Z-stacks with 0.3 μm spacing between slices so that the full 3D structure of SCs could be visualised. The only exception to this was during the time-lapse imaging of the Rab6 to Rab11 transition for which only a single representative Z-plane was imaged in order to minimise bleaching and phototoxicity over the 6-8-hour experiments. The SoftWoRx software was used to deconvolve Z-stacks to improve the signal:noise ratio in images prior to analysis.

### Analysis and parameters

Deconvolved images were analysed in FIJI/ImageJ. To determine the number of compartments marked by a specific *Rab* gene-trap, every compartment was counted that was >1 μm in diameter at its widest point and which displayed fluorescent signal that was distinct from background signal specifically at its limiting membrane. In cases where this was not clear, the transect tool on ImageJ was used to determine whether there was a peak in fluorescent signal at the limiting membrane. To determine the number of DCGs in SCs, the DIC channel was used across all genotypes and was supplemented with the GFP channel in GFP-GPI backgrounds. The DIC channel was also used to assess the morphology of DCGs. The vast majority of DCGs in wildtype backgrounds were uniform, round and each secretory compartment typically contained only a single large central DCG. Compartments which contained abnormal and immature DCGs were therefore defined as having at least one of the following in any Z-plane apart from the first (apical) or last (basal) two in-focus Z-planes: 1) Multiple core ‘fragments’ present within a single compartment; 2) DCGs containing two or more acute external angles; 3) DCGs containing an internal angle greater than 180°.

The presence of Rab6-positive ILVs inside a given compartment was determined by scoring internal fluorescent puncta inside Rab6-marked compartments in *Rab6-CFP* gene trap backgrounds

### Statistical analysis

Statistical significance for all experiments was determined using the non-parametric Kruskal-Wallis test, performed on GraphPad Prism. All graphs displayed in figures show the mean value for each genotype and include error bars representing the standard deviation. n ≈ 30 cells for each genotype, assessed using at least 10 independent AG lobes.

## Acknowledgements

We are extremely grateful to B. Kroeger, who initiated a significant part of the work presented. We thank all the staff at Micron Advanced Bioimaging Unit (supported by Wellcome Strategic Awards 091911/B/10/Z and 107457/Z/15/Z), where imaging was undertaken. We also thank S. Eaton, S. Goodwin, E. Prince and F. Karch, as well as the Bloomington and Vienna *Drosophila* Stock Centres for *Drosophila* stocks. We acknowledge the support of the BBSRC (BB/K017462/1, BB/N016300/1, BB/R004862/1, BB/W00707X/1 and BB/W015455/1) and Cancer Research UK (C19591/A19076, C602/A18974). AW was funded by a studentship from the Biochemical Society’s Krebs Memorial Fund and FC was supported by MINCIENCIAS, Colombia (Call 529). This research was funded in part by the above grants from the BBSRC and the Wellcome Trust. For the purpose of Open Access, the author has applied a CC BY public copyright licence to any Author Accepted Manuscript (AAM) version arising from this submission.

## Supplementary Data

## Supplementary Movies

Movie S1 (related to Fig. 2). Rab1 to Rab6 transition.

Movie S2 (related to Supp. Fig. S2). Fusion of Rab1/Rab6 co-labelled compartments.

Movie S3 (related to Fig. 3). Rab6 to Rab11 transition and DCG biogenesis.

Movie S4 (related to Supp. Fig. S3). Rab6 to Rab11 transition.

---

**Figure S1.**
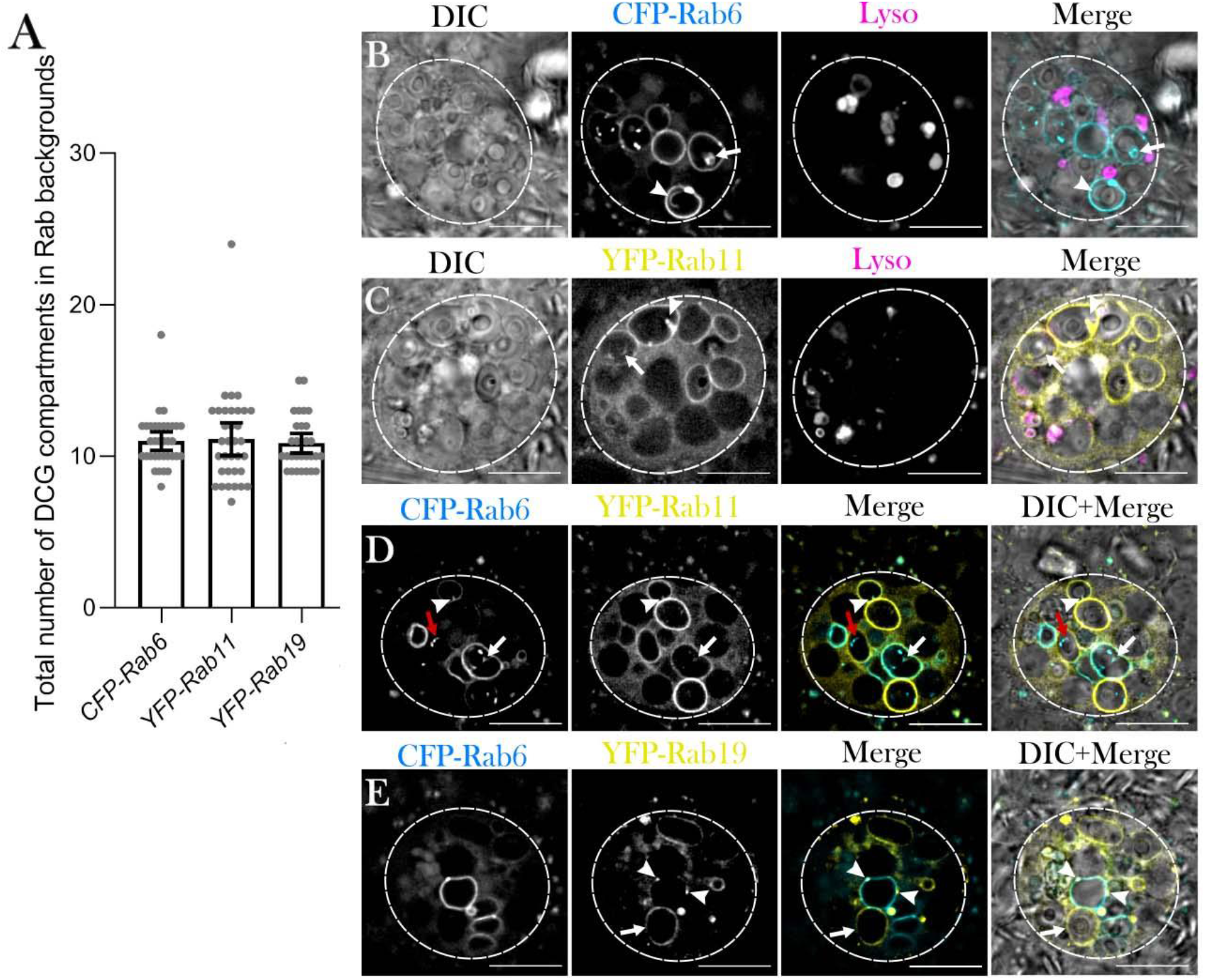
Secretory compartments and ILVs in SCs are marked by several different Rab proteins (related to Fig. 1) (A) Histogram showing the number of DCG-containing compartments in control SCs and in SCs expressing different *Rab* gene-traps, as assessed by DIC microscopy. (B-D) ILVs in SCs associate with DCGs and can form chains extending from the limiting membrane to DCGs. (B) CFP-Rab6-labelled ILVs cluster at the surface of DCGs (arrow) and form chains which extend from the limiting membrane to the DCG boundary (arrowhead). (C) YFP-Rab11-labelled ILVs also cluster at the surface of DCGs (arrow) and form chains which extend from the limiting membrane to the DCG boundary (arrowhead). (D) Clusters of DCG-associated ILVs (white arrow) and ILV chains (arrowhead) can be co-labelled by CFP-Rab6 and YFP-Rab11. Note that there are Rab6-positive puncta inside Rab11-compartments that are Rab6-negative (red arrow). (E) In addition to two or three DCG compartments (eg. arrow), YFP- Rab19 marks microdomains on the surface of CFP-Rab6-labelled compartments, indicated by arrowheads. Data for the histogram were collected from three SCs per gland derived from 10 glands; bars show mean ± SD. Approximate outlines of SCs are marked by dashed circles. Scale bars: 10 µm.

**Figure S2.**
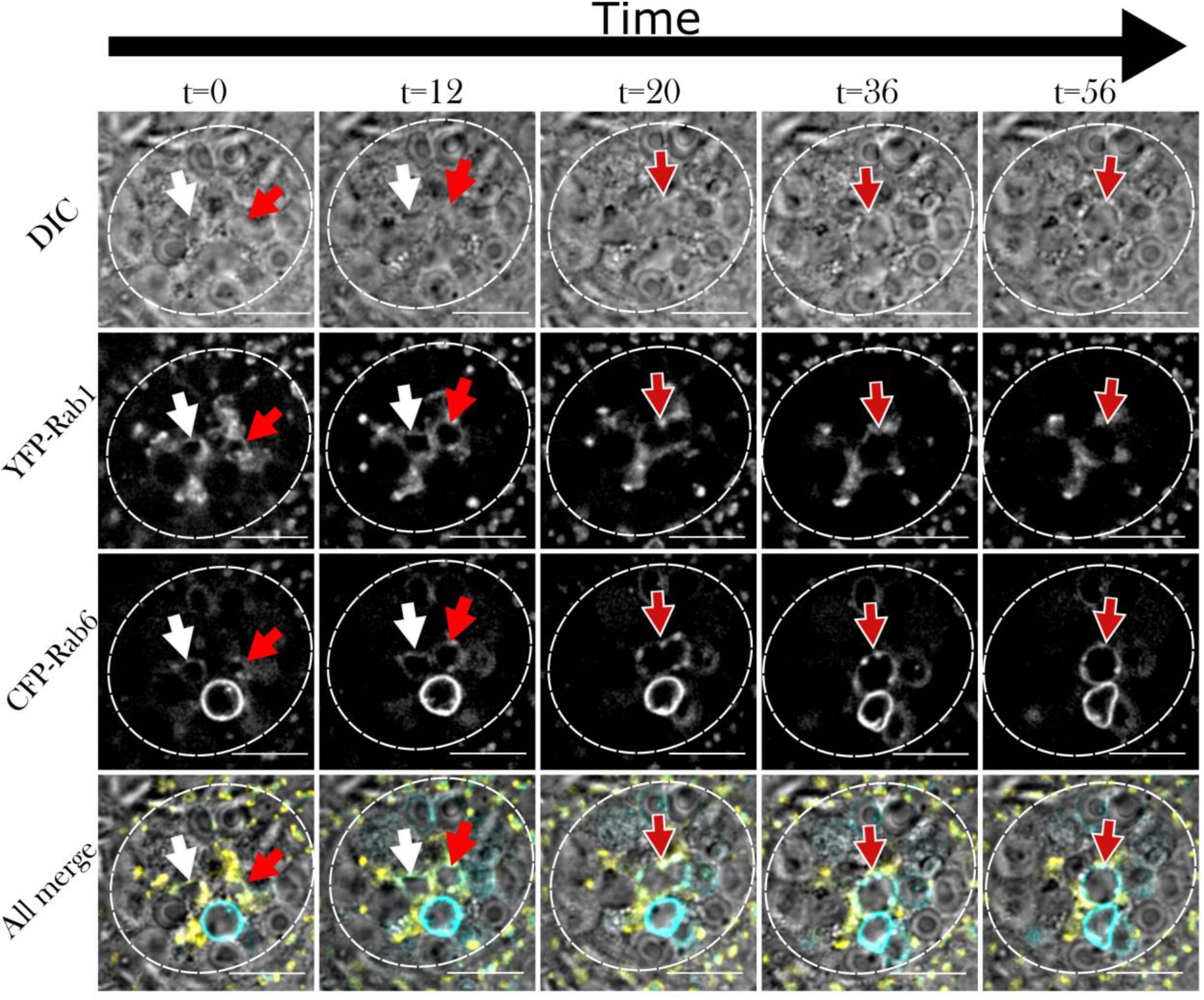
*Trans*-Golgi compartments co-marked by Rab1 and Rab6 can fuse during the Rab1 to Rab6 transition (related to Movie S2) Panel shows *ex vivo* images of a single SC taken at five discrete timepoints with time since first image shown above in minutes. Rows within panel display cellular organisation at each timepoint through DIC imaging (A-E), fluorescent YFP-Rab1 signal (A’-E’), fluorescent CFP-Rab6 signal (A’’-E’’), and combined images displaying all three (A’’’-E’’’). Two compartments marked by both YFP-Rab1 and CFP-Rab6 are marked by a white arrow and a red arrow. After the fusion of these compartments, the combined compartment is denoted by a red arrow with a white outline. (A-A’’’) The two central spherical compartments, which are jointly labelled by YFP- Rab1 and CFP-Rab6, initiated their Rab1 to Rab6 transition and expansion in volume approximately 25 minutes prior to time 0 (see Movie S2). (B-B’’’) After 12 minutes, the two compartments have moved adjacent to each other, as YFP-Rab1 staining gradually diminishes. (C-C’’’) The two compartments fuse to form a single, larger compartment with a distorted shape. (D-D’’’ and E-E’’’) As the time-lapse video continues, the newly formed enlarged compartment regains a spherical shape and loses all YFP-Rab1 labelling. Labelling by CFP-Rab6 continues to increase, resulting in a central, spherical, Rab6-positive compartment which contains no DCG. Approximate outlines of SCs are marked by dashed circles. Scale bars: 10 µm.

**Figure S3.**
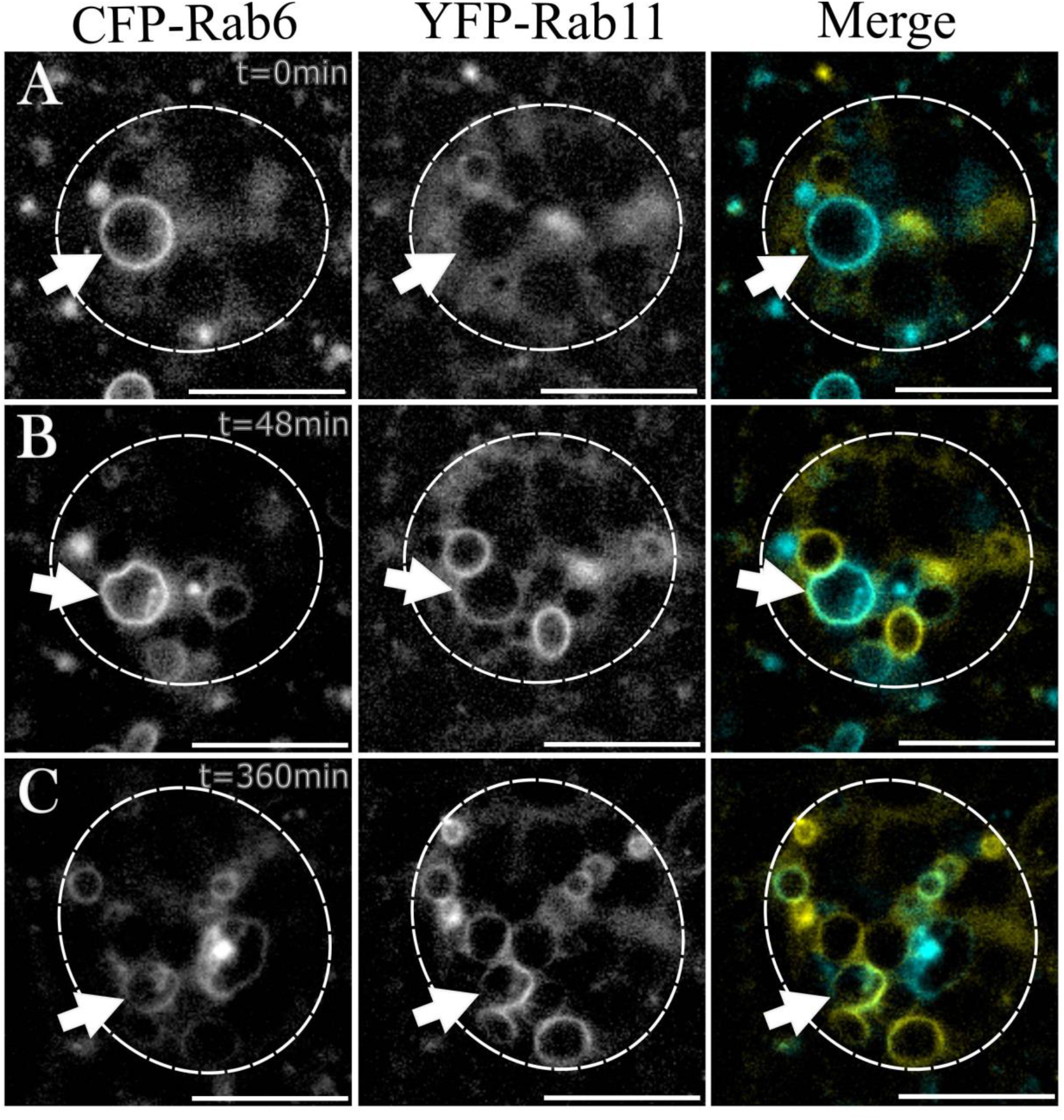
Stages of the Rab6 to Rab11 transition on SC secretory compartments (related to Movie S4) (A-C) Images showing progression of events in the Rab6 to Rab11 transition in SCs expressing the *YFP*-*Rab11* and *CFP-Rab6* gene-traps. White arrows highlight a single maturing compartment at three different timepoints during the transition. (A) Prior to Rab11 accumulation, compartments are marked by Rab6 only and are typically spherical with no ILVs present. (B) Following this, Rab11 starts to accumulate on compartment membranes at low levels. Simultaneously, compartments reduce in size and ILV biogenesis (marked by internal CFP-Rab6 puncta in particular) begins. (C) Over the course of many hours, Rab6 is gradually replaced by Rab11 as the primary marker of these secretory compartments; CFP- Rab6 remains visible on ILVs within compartments. Approximate outlines of SCs are marked by dashed circles. Scale bars: 10 µm.

**Figure S4.**
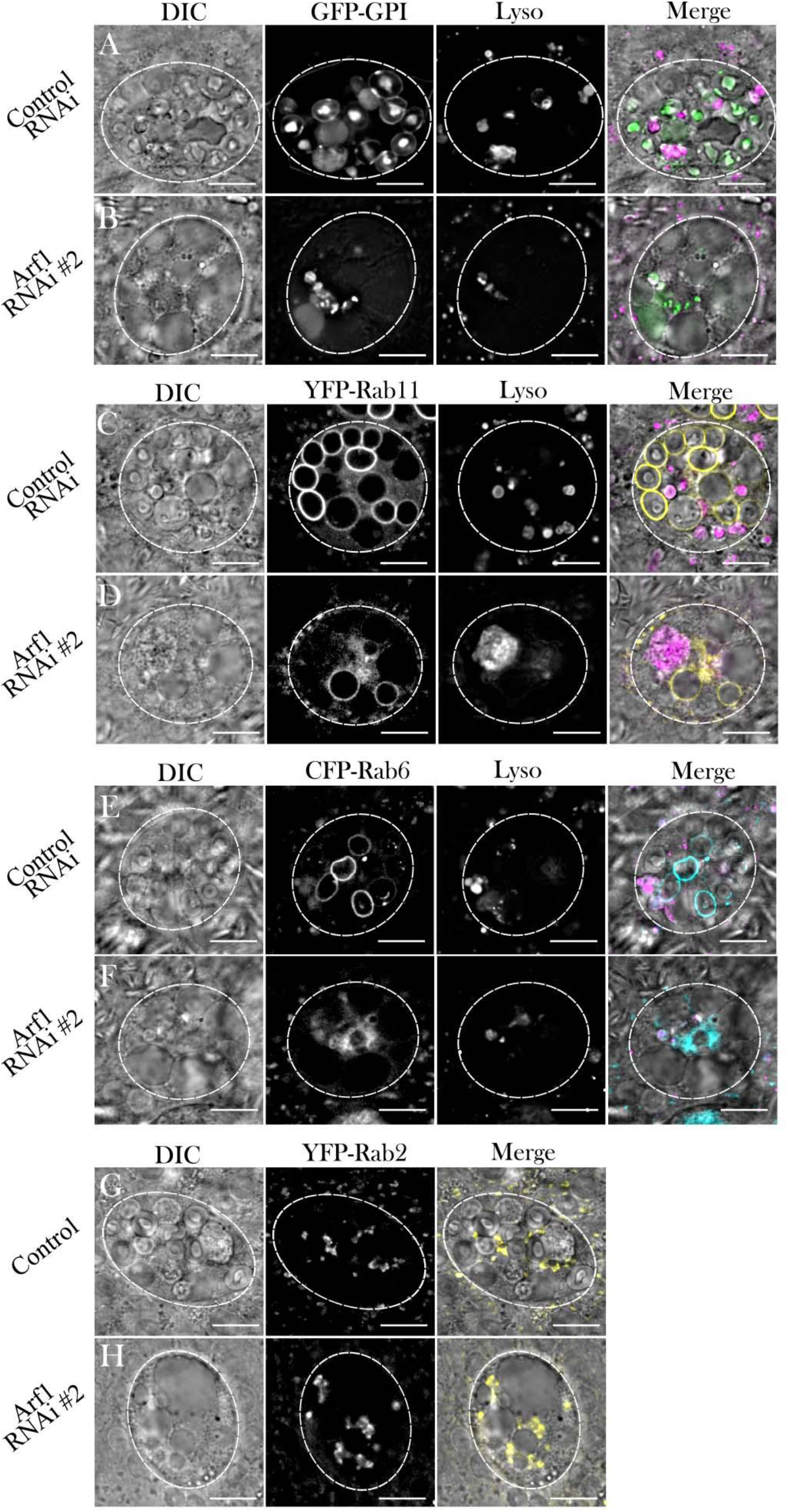
Expression of a second *Arf1* RNAi disrupts DCG biogenesis and Rab6/Rab11 compartment organisation (related to Figs. 4, 5 and 6) (A-F) Representative images of SCs expressing a control RNAi or *Arf1* RNAi #2 in *GFP-GPI* (A, B), *YFP-Rab11* (C, D), and *CFP-Rab6* (E, F) backgrounds. (B) SCs expressing *Arf1* RNAi #2 contain significantly fewer DCGs than controls (A). (D) Rab11-positive compartment organisation is severely disrupted in SCs expressing *Arf1* RNAi #2 versus control (C). (F) Rab6-positive compartment organisation is severely disrupted in SCs expressing *Arf1* RNAi #2 versus control. (G, H) Representative images of SCs expressing the *YFP-Rab2* gene-trap either alone (G) or alongside *Arf1* RNAi #2 (H). When compared to control SCs, the distribution of YFP-Rab2 does not appear to significantly change following *Arf1* knockdown and YFP-Rab2 does not mark any large non-acidic compartments. Approximate outlines of SCs are marked by dashed circles. Scale bars: 10 µm.

**Figure S5.**
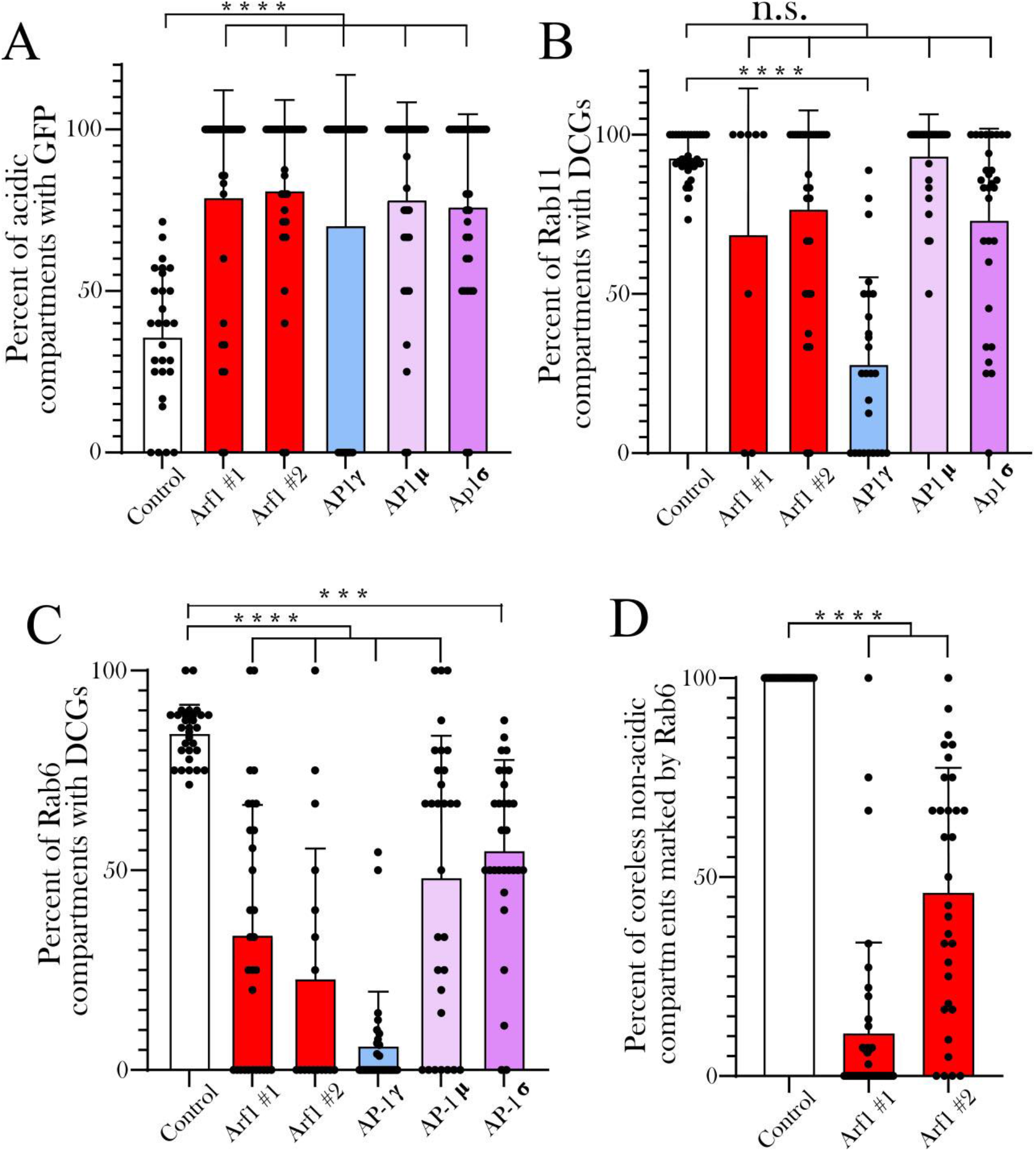
Knockdown of *Arf1* and *AP-1* subunits affects GFP-GPI trafficking to acidic compartments and the identity of non-acidic compartments in SCs (related to Figs 4, 5 and 6) (A) Histogram showing the proportion of acidic-compartments containing unquenched GFP in control SCs or following knockdown of *Arf1* or *AP-1* subunits. (B) Histogram showing the proportion of Rab11-positive compartments which contain DCGs in these different genotypes. Note that despite the significant decrease in DCGs seen in knockdowns (Figs. 4F, 5G), the proportion of Rab11-positive compartments which contained DCGs was not significantly affected by any knockdown other than *AP-1γ*, although most of these DCGs were irregularly shaped (Fig. 5H). Therefore, transition to Rab11 identity typically appears to be associated with subsequent DCG formation. (C) Histogram showing the proportion of Rab6- positive compartments which contain DCGs in these different genotypes. (D) Bar chart showing that knockdown of *Arf1* affects the Rab6-positive identity of large non-acidic compartments that do not contain DCGs. P<0.001: *** P<0.0001: ****.

**Figure S6.**
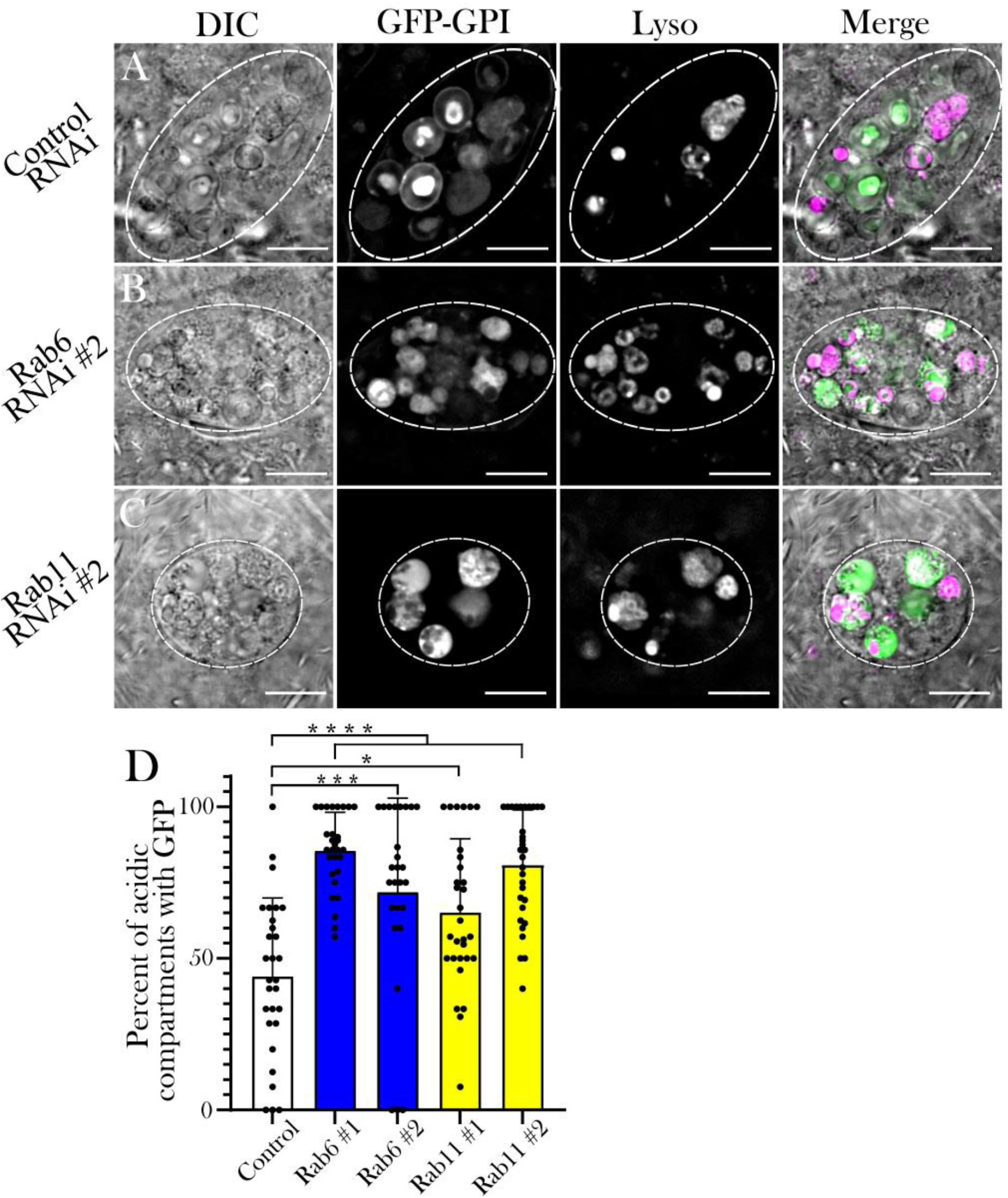
Expression of a second *Rab6* and *Rab11* RNAi disrupts DCG biogenesis (related to Fig. 7) (A-C) Representative images of SCs expressing the DCG marker GFP-GPI with a control RNAi (A) or RNAis (#2) targeting *Rab6* (B) or *Rab11* (C). Cellular organisation is assessed through DIC imaging, GFP-GPI fluorescence and Lysotracker Red fluorescence, as well as a merged image for each cell. In both the *Rab6* and the *Rab11* knockdown, significantly fewer DCGs are present and marked by GFP-GPI. (D) Histogram showing the proportion of acidic compartments containing unquenched GFP in these different genotypes. Approximate outlines of SCs are marked by dashed circles. Scale bars: 10 µm. P<0.05: * P<0.01: ** P<0.001: *** P<0.0001: ****.

**Figure S7.**
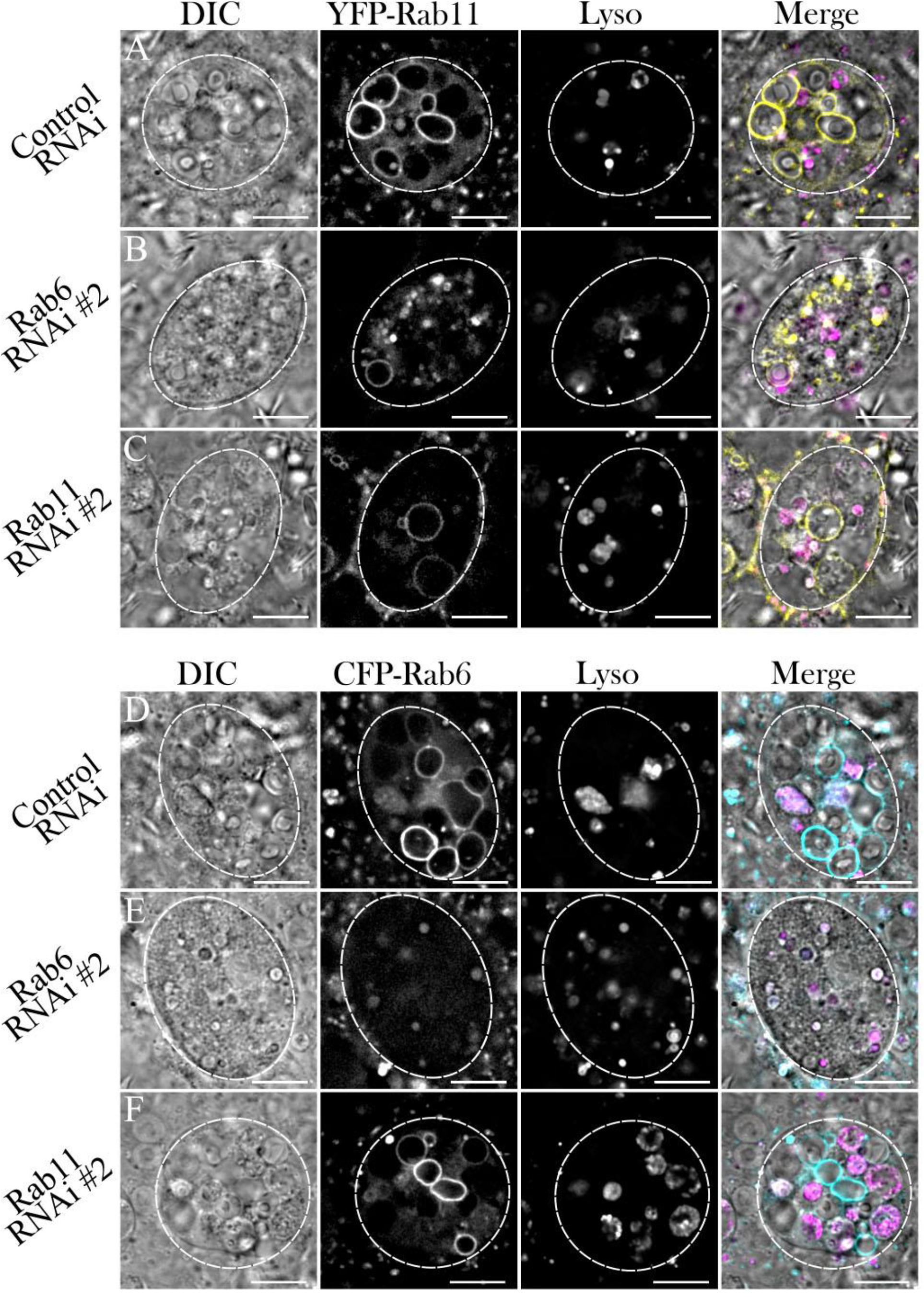
Expression of a second *Rab6* and *Rab11* RNAi affects Rab6- and Rab11-positive compartment organisation (related to Fig. 8) (A-C) Representative images of SCs expressing the *YFP*-*Rab11* gene trap together with a control RNAi (A) or RNAis targeting *Rab6* (B) or *Rab11* (C). (D-F) Representative images of SCs expressing the *CFP*-*Rab6* gene trap together with a control RNAi (D) or RNAis targeting *Rab6* (E) or *Rab11* (F). Cellular organisation in all genotypes is assessed through DIC imaging, gene trap fluorescence and Lysotracker Red fluorescence, as well as a merged image for each cell. Note that some Rab11 gene trap fluorescence is still visible even after knockdown of *Rab11* (C). Approximate outlines of SCs are marked by dashed circles. Scale bars: 10 µm.

